# Seed tuber imprinting shapes the next-generation potato microbiome

**DOI:** 10.1101/2023.07.27.549298

**Authors:** Yang Song, Jelle Spooren, Casper D. Jongekrijg, Ellen H.H. Manders, Ronnie de Jonge, Corné M.J. Pieterse, Peter A.H.M. Bakker, Roeland L. Berendsen

## Abstract

Potato seed tubers are colonized and inhabited by soil-borne microbes, some of which can positively or negatively impact the performance of the emerging daughter plant in the next season. In this study, we investigated the intergenerational inheritance of microbiota from seed tubers to next-season daughter plants by amplicon sequencing of bacterial and fungal microbiota associated with tubers and roots of two seed potato genotypes produced in six different fields. We observed that field of production and potato genotype significantly affected the seed tuber microbiome composition and that these differences persisted during winter storage of the seed tubers. When seed tubers from different production fields were planted in a single trial field, the microbiomes of daughter tubers and roots of the emerging plants could still be distinguished according to the field of origin of the seed tuber. Remarkably, we found little evidence of direct vertical inheritance of field-unique microbes from the seed tuber to the daughter tubers or roots. Hence, we hypothesize that this intergenerational “memory” is imprinted in the seed tuber, resulting in differential microbiome assembly strategies depending on the field of production of the seed tuber.

## Introduction

The microbial community associated with a plant, referred to as the plant microbiome, can significantly influence plant performance. The complex plant microbiome includes microbes that are plant pathogens but also plant beneficial microbes that support plant growth by mobilizing scarce nutrients from the soil or protect the plant against pathogens ^1, 2, 3^. The plant microbiome significantly expands the genomic potential of its host and is often referred to as the host’s “second genome” ^1, 2, 4, 5, 6, 7^.

Potato is the 3rd most important crop for human consumption, with an annual global harvest of approximately 375 million tons. Additionally, it is a key crop that is essential for global food security and a source of raw materials for industry (www.fao.org, 2017). Potato is a space-efficient crop, yielding five times more consumable weight per hectare than rice and wheat. As global demand for potato increases, the UN-FAO identified it as a crop with great potential to become a game changer for global food security ^8, 9^.

Potatoes are commonly propagated vegetatively by transplanting seed tubers from one field to the next ^10^. As potato tubers develop underground, they closely interact with the dense and diverse microbial communities in soil ^11^. Studies demonstrated that the potato tuber microbiome can have a profound impact on plant health and productivity ^12, 13^. Potato is sensitive to a wide range of plant pathogens ^14, 15^, but it also hosts beneficial microbes that can promote plant growth ^12, 13, 16, 17, 18^.

A batch of seed potatoes is of high vitality if it manifests in a large canopy and exhibits homogeneous growth in the early stages of its development. Seed tubers of the same potato genotype that were produced in different production fields can display significant differences in their vitality, resulting in differences in growth of the emerging potato plants ^19, 20, 21^. This may be caused by local environmental factors in the fields of production that confer changes in tuber physiology, but also the seed tuber microbiome likely impacts the vitality of the outgrowing potato crop. Many potato pathogens can be seed tuber-borne ^15, 22^. Field experiments in which seed tubers were treated with beneficial bacteria show that the applied microbes colonize the roots of plants that develop from the treated tubers ^23, 24^. Such findings suggest that seed tubers can be an important inoculum source of microbes for the potato plants that emerge from them and that potato plants may inherit at least part of their microbiome from the seed tuber. However, there is limited information available on tuber-borne transmission of microbes from one potato generation to the next. To gain insight into intergenerational inheritance of the potato microbiome, we investigated whether the field of production of potato seed tubers has an impact on the microbiomes of tubers and roots of plants emerging from these seed tubers when planted together in a single trial field.

## Results

### Effect of potato genotype, production field, and storage on the tuber microbiome

In the autumn of 2018, seed tubers of two potato varieties, *Colomba* (hereafter Variety A) and *Innovator* (hereafter Variety B), were harvested from 3 fields of production for Variety A and 3 different fields for Variety B (Fig. 1a; Fig. S1a-b). To investigate the influence of plant genotype and field of production on the tuber-associated microbiome, we isolated microbial DNA from 4 replicate samples per field, each replicate containing peels of 6 tubers. Subsequently, we sequenced 16S rRNA gene and ITS amplicons to profile the bacterial and fungal communities, respectively. Principal coordinates analysis (PCoA) and permutational multivariate analysis of variance (PERMANOVA), revealed that both the bacterial and the fungal microbiome on the tuber is determined primarily by the field in which the potato was produced (Fig. 1b-d; Fig. S2a-c; Table S1; Table S2). The production field significantly (*P* = 0.001) affected the tuber peel microbiome and accounted for up to 64% of the variation in the bacterial community (R^2^ = 0.64, Table S1) and 55% of the variation in the fungal community (R^2^ = 0.55, Table S2). In addition, the potato variety significantly (*P* = 0.001) affected tuber microbiome composition, explaining 18% (R^2^ = 0.18) and 17% (R^2^ = 0.17) of the variation in bacterial and fungal community composition, respectively (Fig. 1b-d; Fig. S2a-c; Table S1; Table S2).

It is common agricultural practice to store seed tubers over the winter prior to planting in spring. To study the effects of cold storage on tuber microbiomes, the above-mentioned seed tubers had remained in cold storage at 4°C in the dark for 7 months (Fig. 1a). These so-called post-storage seed tubers were then processed in the same manner as the seed tuber samples, after which the bacterial and fungal microbial communities were profiled by amplicon sequencing. Although there were significant changes in the composition of the bacterial (*P* = 0.001) and fungal (*P* = 0.019) microbiome before and after storage of the tubers (Fig. S3a and c, Table S3), post-storage seed tubers clustered closely with those of the pre-storage seed tubers from the same field of production (Fig. S3b and d). Notably, tubers from different fields of production maintained their distinct microbial community patterns even after 7 months of cold storage (Fig. 1e-g; Fig. S2d-f; Table S1; Table S2). On post-storage seed tubers, the production field accounted for up to 57% of the variation in the bacterial community (R^2^ = 0.57, Table S1) and 46% of the variation in the fungal community (R^2^ = 0.46, Table S2).

**Fig. 1.**
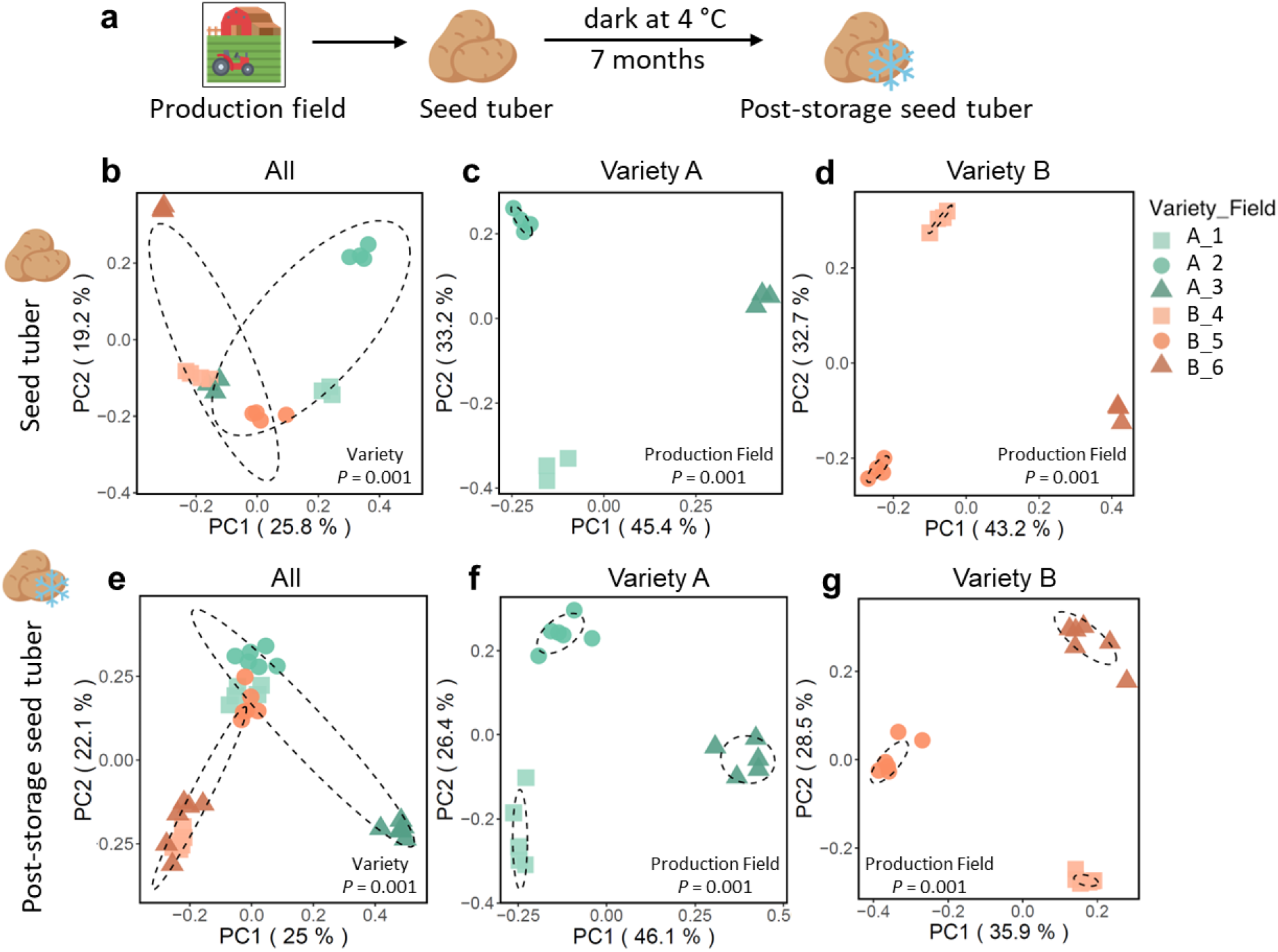
Bacterial community composition of seed tuber and post-storage seed tuber samples. **a** Graphic representation of the sampling strategy. Seed tubers of two potato varieties (A and B) were harvested from 3 fields of production per variety and sampled for microbiome analysis before and after a 7-month cold storage period. Principle component analysis (PCoA) of 16S amplicon sequencing data representing bacterial communities on **b)** seed tubers of Variety A and B, **c)** seed tubers of Variety A only, or **d)** seed tubers of Variety B only, **e)** post-storage seed tubers of Variety A and B, **f)** post-storage seed tubers of Variety A only, and **g)** post-storage seed tubers of Variety B only. Each symbol represents the bacterial community of one replicate potato peel sample. Each sample consists of a pool of potato peels collected from 6 seed tubers. For each variety, 4 replicate of seed tuber samples and 6 replicate of post-storage seed tuber samples were collected from each of the 3 fields of production. Green symbols represent Variety A and orange symbols represent Variety B. Different shapes within a same color represent distinct production fields. The *P* from PERMANOVA is shown in each PCoA plot. Each ellipse represents a 68% confidence region and depicts the spread of data points within each group.

### Seed tubers, roots of emerging plants, and daughter tubers harbor distinct microbiomes

Seed tubers of Variety A and B of the above-mentioned 6 production fields were subsequently planted in a single trial field near Veenklooster, the Netherlands, in the spring of 2019 (Fig. S1a). The emerging plants from these seed tubers were cultivated for three months after which roots and daughter tubers were harvested (Fig. 2a). The microbiome composition of these potato samples was analyzed by sequencing both 16S rRNA gene and ITS amplicons. Using PCoA, we observed that the bacterial community composition of both roots and tubers harvested in 2019 from the trial field clearly separated (*P* = 0.001) from the seed tuber samples harvested from the production fields in 2018 (Fig. 2b; Table S4). In addition, the bacterial communities found on the roots are distinct from those on daughter tubers, indicating that these two belowground potato organs harbor distinct bacterial microbiomes within one field (*P* = 0.001, Fig. 2b; Table S4). A similar separation was observed for the fungal communities on seed tubers, roots, and daughter tubers (Fig. S4a, Table S4).

We then focused on shared bacterial amplicon sequence variants (ASVs) between the microbiomes of seed tuber, daughter tuber, and root samples (Fig. 2c). A total of 3986, 9205, and 11622 unique bacterial ASVs were detected in seed tuber, daughter tuber, and root samples, respectively. Whereas 86% ((6830+1050)/9205) of the bacterial ASVs on the daughter tubers were shared with those on roots of the potato plants in the same trial field, only 13% ((1050+156)/9205) and 12% ((1050+393)/11622) of the ASVs on the daughter tubers and roots, respectively, were also detected on the seed tuber (Fig. 2c). Analysis of the fungal microbial communities showed similar results with 84% ((758+182)/1117) of the fungal ASVs from daughter tubers shared with those on roots, while only 18% ((182+22)/1117) and 16% ((182+37)/1405) of the ASVs detected on the daughter tubers and roots, respectively, were also detected on the seed tubers (Fig. S4b). This suggests that the majority of microbes on potato daughter tubers and roots are not inherited from the seed tubers but originate from the trial field.

The most abundant bacterial phyla in the microbiomes of all tuber and root samples were the *Proteobacteria, Actinobacteria, Bacteroidetes* and *Firmicutes*. Whereas *Bacteroidetes* were relatively abundant in samples from plants in the trial field, *Firmicutes* had relatively low abundance in samples from this field, especially on the daughter tubers. On those daughter tuber samples *Bacteroidetes* were relatively more abundant, whereas *Actinobacteria, Firmicutes*, and *Planctomycetes* had higher relative abundance on the roots of the potato plants in the same field (Fig. 2d).

**Fig. 2.**
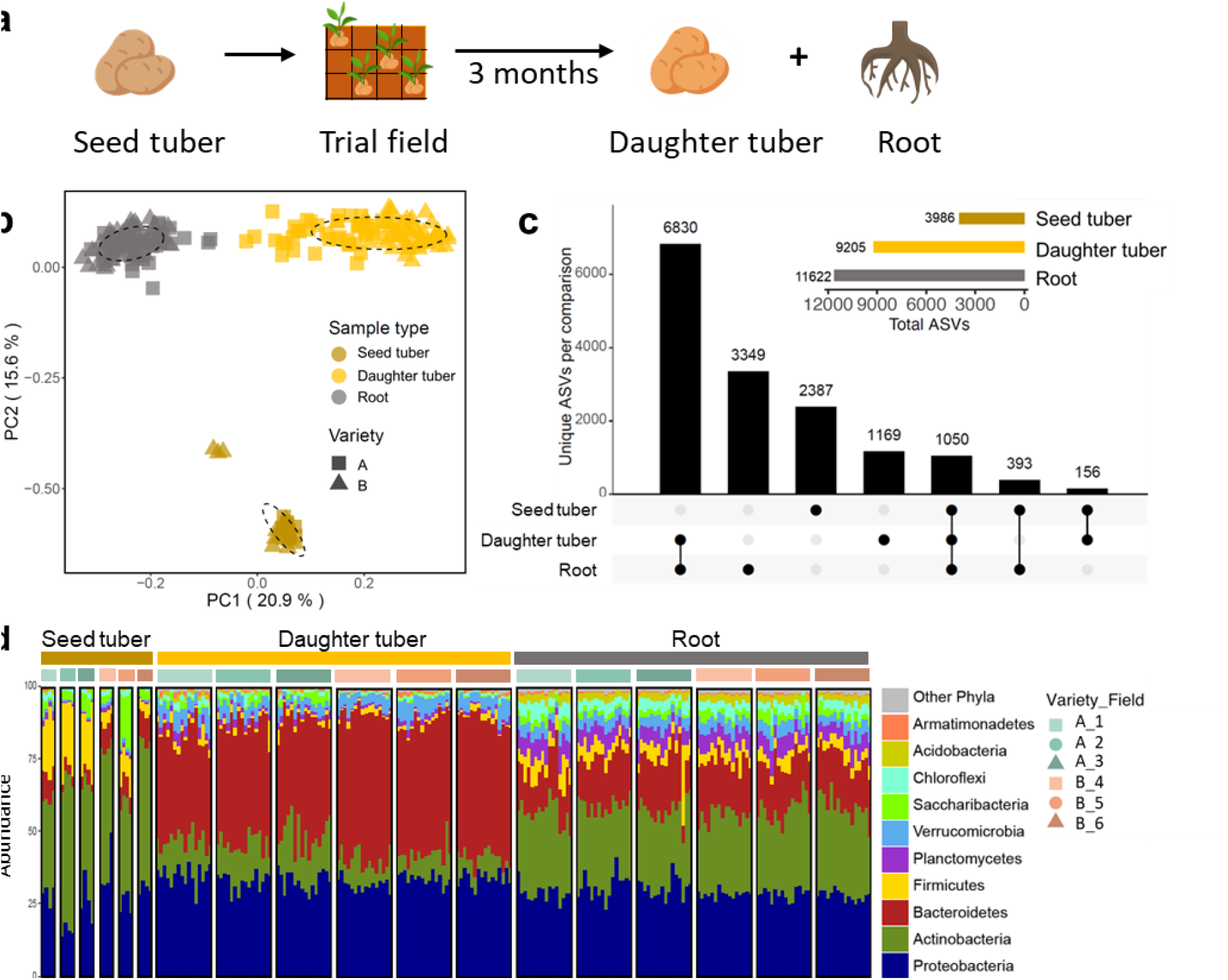
Analysis of bacterial communities on seed tubers from different production fields and their roots and daughter tubers the Veenklooster trial field. **a** Graphic representation of the experimental design. Seed tubers from 3 different production fields of each variety (*n =* 2) were planted together in a single trial field in Veenklooster (Fig. S1). For seed tubers from each production field, 4 replicate plots were randomly distributed across this trial field. Roots and daughter tubers from the emerging plants were harvested for microbiome analysis. **b** PCoA of potato-associated bacterial communities of seed tubers, daughter tubers, and roots. Square symbols represent Variety A and triangle symbols represent Variety B. Colors represent different sample types. Each ellipse represents a 68% confidence region and depicts the spread of data points within each group. **c** UpSet plot showing the number of bacterial ASVs that are shared between or are unique for seed tubers, daughter tubers and roots of both varieties combined. **d** Stacked bar chart of the taxonomic composition of bacterial communities of different sample types aggregated at the phylum level. Each stacked column represents an independent sample (n = 216). Different colors within a column represent different phyla. Only the top 10 most-abundant phyla were colored individually, all the rest are colored in gray and listed as “Other phyla”. Samples are clustered by sample type and production field, which is shown by the colored bar on top of the stacked bar chart.

### Origin of seed tubers affects the root and tuber microbiomes of emerging plants

Within the trial field, the bacterial and fungal microbial communities of both potato roots and daughter tubers were significantly (*P* = 0.001) affected by potato genotype (Fig. 3a-b, Fig. S2g and j, Table S1, Table S2). The effect size of potato genotype was larger for the tuber samples (R^2^ = 0.08) than for the root samples (R^2^ = 0.03). Interestingly, also the field of production of the seed tubers had a significant effect on microbiome composition of daughter tubers (*P* = 0.001, R^2^ = 0.07) and roots (*P* = 0.001, R^2^ = 0.08) of the plants emerging from these seed tubers (Fig. 3c-f, Fig S2g-l, Table S1, Table S2). Thus, the impact of the production field stretches across a generation and influences microbiome assembly on the roots and tubers of the daughter plants emerging from the seed tubers in the subsequent growing season.

**Fig. 3.**
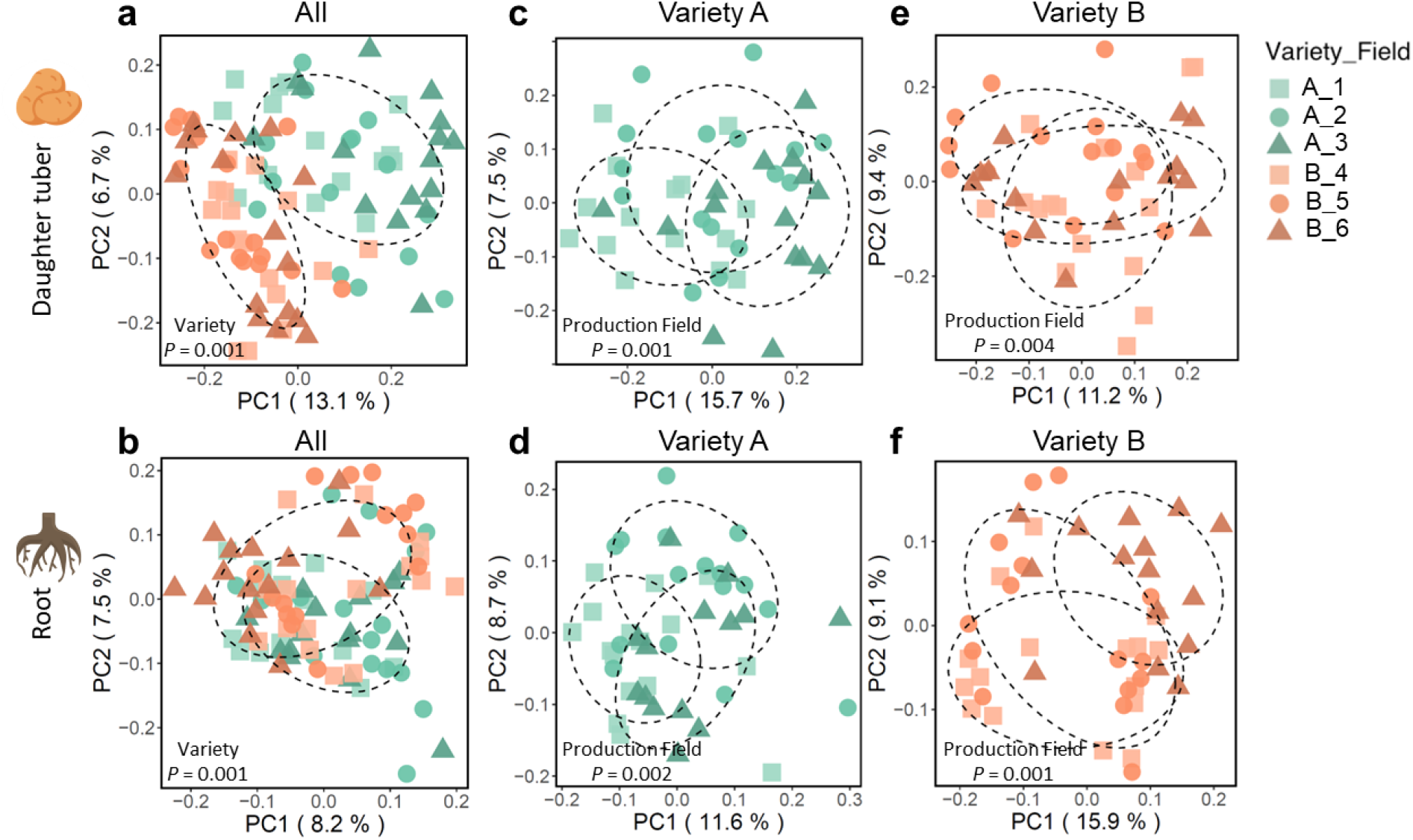
Bacterial community composition of daughter tubers and roots. PCoA of 16S rRNA amplicon sequencing data of **a)** daughter tubers of Variety A and B, **b)** roots of Variety A and B, **c)** daughter tubers of Variety A only, or **d)** roots of Variety A only, **e)** daughter tubers of Variety B only and **f)** roots of Variety B only. Each symbol represents the bacterial community of one replicate potato peel sample. Each daughter tuber sample consisted of a pool of potato peels collected from 6 daughter tubers of one plant. Each root sample is a subset of the whole root of the same plant from which the daughter tubers were sampled. For each variety, 4 replicate samples were collected from each of the 4 randomly distributed replicate plots. Green symbols represent Variety A and orange symbols represent Variety B. Different shapes within a same color represent different production fields. The *P* from PERMANOVA is shown in each PCoA plot. Each ellipse represents a 68% confidence region and depicts the spread of data points within each group.

### Inheritance of field-unique ASVs in daughter tubers and roots

We hypothesized that the intergenerational influence of the seed tuber production field on the microbiome of roots and daughter tubers is the result of vertical, seed tuber-mediated transmission of field-unique microbes from one generation of potatoes to the next. To be able to track the vertical inheritance of field-unique microbes from seed tubers to the emerging plants, we focused on Variety A seed tubers from Field 1 and identified bacterial and fungal ASVs that were uniquely detected in seed tuber samples from Field 1. We observed that 50.6% of the bacterial ASVs on seed tubers from Field 1 were not detected on seed tubers from Field 2 and 3 and defined these 952 ASVs as Field-1-unique on seed tubers (Fig. S5a). With the same definition, we identified 1451 bacterial ASVs (29.7% of total daughter tuber ASVs) as Field-1-unique on daughter tubers that originate from Field-1 seed tubers and 1244 bacterial ASVs (20.2% of total root ASVs) as Field-1-unique on roots that originate from Field-1 seed tubers (Fig. S5c-d). An additional 54, 132 and 137 fungal ASVs were defined as Field-1-unique on seed tubers, daughter tubers, and roots, respectively (Fig. S5e-h).

We subsequently investigated whether the Field-1-unique ASVs were transmitted to the roots and daughter tubers of the plants emerging from these Field-1 seed tubers. To our surprise, the results did not support our original hypothesis, and instead, we found only a very small overlap between Field-1-unique ASVs of seed tubers and daughter tubers and roots derived from these Field-1 seed tubers (Fig. 4a-b and d-e, Fig. S6a-b and d-e). Namely, only 24 bacterial and 3 fungal Field-1-unique ASVs were shared between seed tubers and the emerging daughter tubers. Similarly, only 26 bacterial and 1 fungal Field-1-unique ASVs were shared between seed tubers and the roots of emerging plants (Fig. 4a-b, Fig. S6a-b). Moreover, these ASVs were lowly abundant in daughter tuber (Bacteria: 0.1%, fungi: 0.3%) and root (Bacteria: 0.05%, fungi: 0.09%) microbial communities (Fig. 4g-h, Fig. S6g-h). Thus, although we can distinguish ASVs unique to the field of production of the seed tuber on the next season daughter tubers and roots, the large majority of the field-unique ASVs in the daughter generation cannot be immediately traced back to the seed tuber.

When looking into the entire microbial community on seed tubers instead of only the field-unique ones, we found that 83% (1556/1882, Fig. 4d) and 78% (1472/1882, Fig. 4e) of the seed tuber bacterial ASVs were lost during vertical transmission to daughter tubers and roots, respectively. Furthermore, 77.2% of the daughter tuber (Fig. 4g) and 74.5% (Fig. 4h) of the root bacterial communities were acquired from the environment during the 3 months of growth in the trial field. Around a quarter of daughter tuber (22.8%, Fig. 4g) and root (25.6%, Fig. 4h) microbial communities were shared with those on the peel of the seed tuber. However, since these ASVs were not Field 1-unique, it cannot be verified to what extent they are inherited from the seed tuber or simply common in different fields. Similar results were observed for the fungal communities on the daughter tubers and roots from Field 1 (Fig. S6). These results indicate that even though the field-unique ASVs were rarely inherited cross generations, we did observe vertical inheritance for other ASVs from seed tubers to daughter tubers and roots. However, the majority of the microbial population in daughter tubers and roots were acquired from the environment where they were formed.

To investigate whether cold storage would already lead to the depletion of the above defined field-unique seed tuber microbes pre-planting, we examined the occurrence of ASVs on the post-storage seed tubers from Field 1. These post-storage seed tubers were stored under cold and dark condition much longer than common practice, thus used as an extreme case to study the influence of storage on field-unique seed tuber microbes. We found that 66% (1051/1593) of the total bacterial ASVs detected on the post-storage seed tubers were also detected on the pre-storage seed tubers from the same field (Fig. 4c and f) and that these ASVs represent 91.8% of the bacterial community (Fig. 4i). These results indicated that the large majority of the seed tuber bacterial community persists during cold storage. Moreover, a large part of the field-unique ASVs were maintained over the storage period (Fig. 4f and i, “Unique-Unique”). Similar results were observed for fungal communities on the seed tuber and post-storage seed tubers from Field 1 (Fig. S6f and i).

**Fig. 4.**
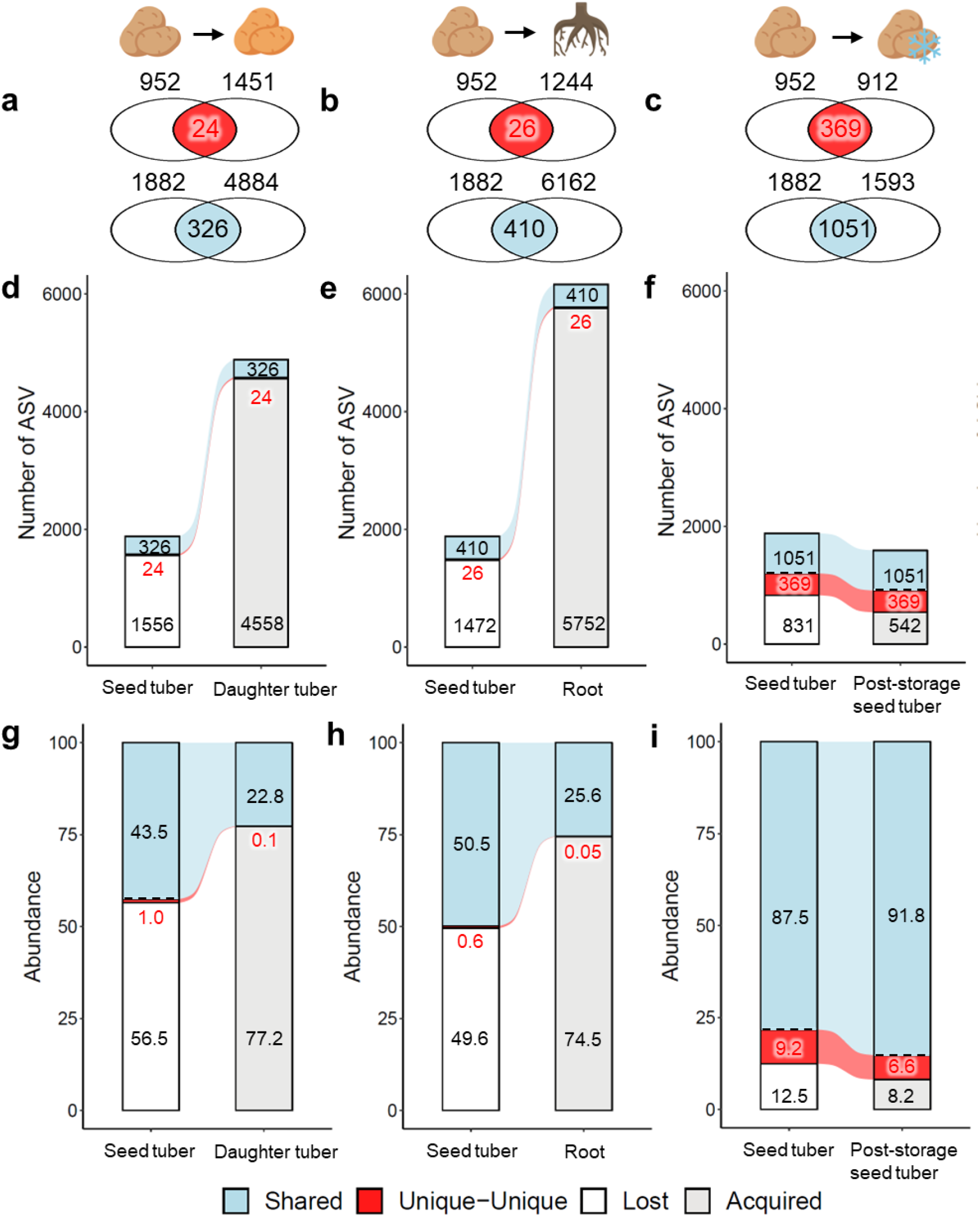
Comparison of bacterial ASVs on daughter tubers, roots, post-storage seed tubers and seed tubers of Variety A originating from Field 1. Venn diagrams showing the overlap between **a)** seed tubers and daughter tubers, **b)** seed tubers and roots, **c)** seed tubers and post-storage seed tubers of Field-1-unique bacterial ASVs (in red) or all bacterial ASVs (in blue). Sankey diagram of bacterial ASVs transferred from seed tubers to **d, g)** daughter tubers and **e, h)** roots that emerged from the seed tubers; and **f, i)** post-storage seed tubers. “Shared” in blue represents ASVs detected on both sample types. “Unique-Unique” in red represents the overlap of Field-1-unique ASVs on both sample types. The “Unique-Unique” in red is included in the “Shared” in blue. “Lost” in white represents ASVs lost from the seed tuber during vertical transmission. “Acquired” in light grey represents ASVs not transmitted from seed tubers but acquired from the environment. In **a-f)**, numbers in the bars indicate the number of ASVs in each category mentioned above. In **g-i)**, numbers in the bars indicate the accumulative relative abundance of ASVs in each category mentioned above.

### Tracking the microbial transmission from different seed tuber compartments to sprouts

Even though the microbiomes on daughter tubers and roots of next-season potato plants could be distinguished based on the field of production of the seed tuber, we found little evidence for direct vertical transmission of microbes from the peel of the seed tuber to the peel of the tubers or roots on the daughter plants. This could mean that: 1) potato daughter plants do not inherit their microbiome from the peel but other compartments of the seed tuber; or 2) vertical transmission is apparent only during early stages of plant development after which transmitted microbes are replaced by members from the trial field resident microbiome. To gain further insight into the potential of vertical microbiome transmission from seed tubers to next-generation daughter plants, we investigated the contribution of different seed tuber compartments (namely peel, eye, heel end, flesh, and adhering soil, Fig. S1c) in shaping the potato sprout microbiome. We made use of material from a parallel study in which we harvested tubers from 6 potato varieties produced in 25 distinct fields of production (Variety A from 5, Variety B from 5, *Festien* (Variety C) from 3, *Challenger* (Variety D) from 5, *Sagitta* (Variety E) from 5, and *Seresta* (Variety F) from 2 fields, respectively; Fig. S1b). Samples from 50 seed tubers were pooled into a single sample per compartment per field and thus a total of 1250 (50 x 25) tubers were sampled from these 25 fields. DNA was isolated and bacterial and fungal microbiome composition was determined through 16S rRNA gene and ITS amplicon sequencing.

Again we found that potato genotype significantly influenced the composition of bacterial and fungal communities in the distinct seed tuber compartments (Fig. 5a, Fig. S7a, Table S1, Table S2). Moreover, we found that each distinct tuber compartment harbored a bacterial community that is significantly different (*P* < 0.001) from the communities in the other compartments (Fig. 5b-c, Table S5), with the exception of the pairwise comparisons between eye and peel (*P* = 0.143) and between eye and heel end (*P* =0.061). The richness of the bacterial communities decreased from the outside of the potato to the inside, with highest diversity in the potato-adhering soil and increasingly lower diversity in respectively the potato peel, heel end, eye, and flesh compartments (Fig. S8a). At phylum level, *Bacteroidetes* and *Proteobacteria* have a higher relative abundance in the heel end compartments compared to the other 4 tuber compartments (Fig. 5c).

Similar to the bacterial communities, fungal communities found in distinct compartments were significantly different from each other (*P* < 0.001, Fig. S7b, Table S6), except for the eye and peel compartments which harbored nearly identical fungal communities (*P* = 0.83, Table S6). The highest richness for fungal communities was observed in adhering soil samples; however, diversity did not differ significantly between the other compartments (Fig. S8b). At family level, *Cladosporiaceae* was most abundant in the adhering soil, whereas *Plectosphaerellaceae* was relatively more abundant in the heel ends (Fig. S7c).

**Fig. 5.**
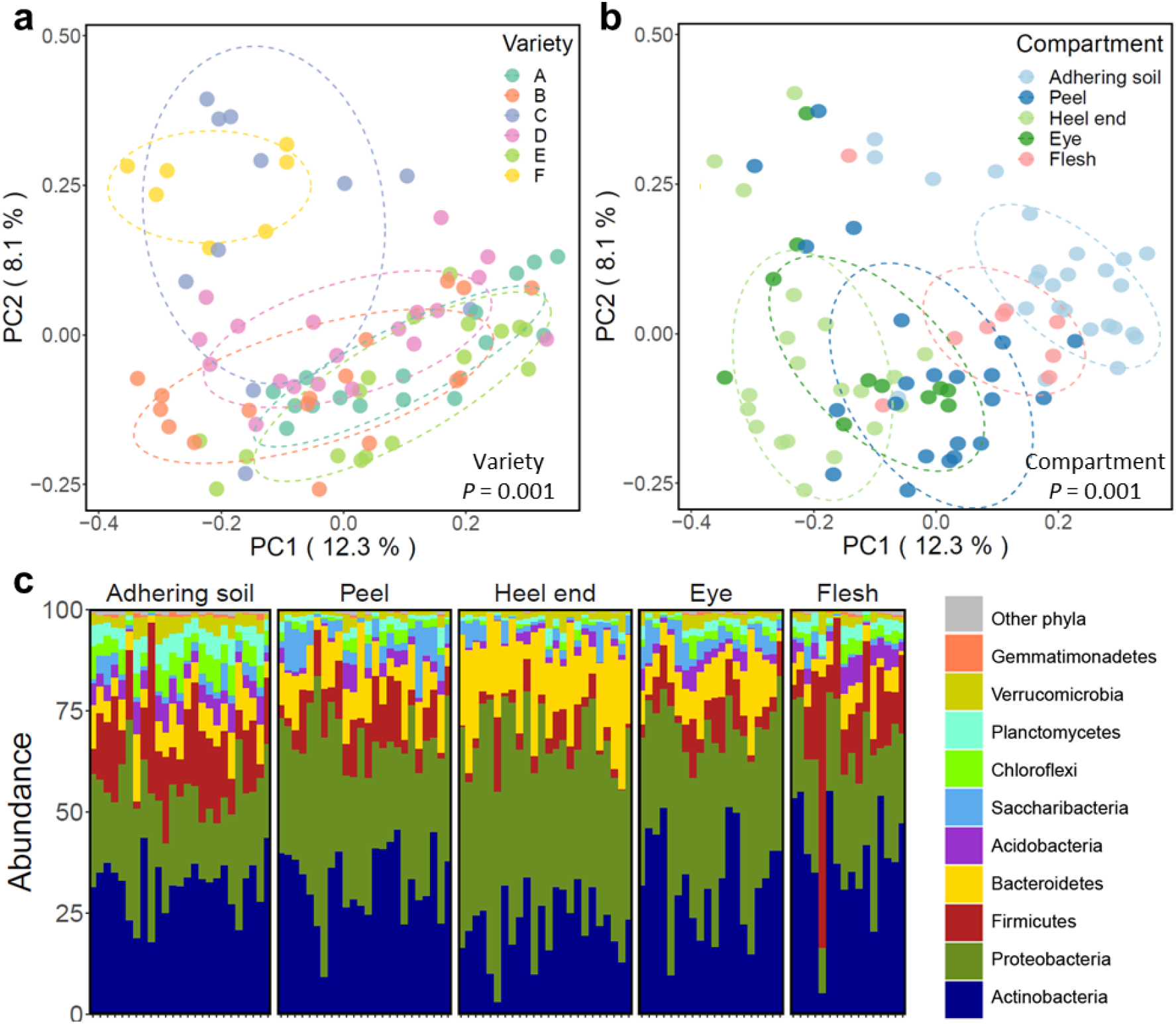
Distinct compartments on the potato tuber harbor distinct microbial communities. PCoA of the potato tuber-associated bacterial community based on 16S rRNA gene amplicon sequencing and colored by **a**) potato varieties (A, B, C, D, E and F, respectively) and **b**) potato tuber compartments (adhering soil, peel, heel end, eye and flesh, respectively). *P* as determined by PERMANOVA is shown in each PCoA plot. Each ellipse represents a 68% confidence region and depicts the spread of data points within each group. **c** Bar plot showing the phylogenetic composition of the bacterial community. Only the top 10 most abundant phyla are colored individually, the other phyla are shown together in grey. Each sample was isolated from the pooled compartments from 50 seed tubers per field.

The spout is the first daughter tissue to emerge from the seed potato, and thus the most likely tissue for vertical transmission of microbiota. To investigate vertical transmission of microbes from the seed tuber to the emerging plant, seed tubers of all 6 varieties and from 2 fields per variety sprouted on Petri dishes for 7 days. Subsequently, we isolated microbial DNA of sprouts of 5 replicate tubers per field and analyzed microbiome composition of the samples through 16S rRNA gene and ITS amplicon sequencing. The bacterial community composition of sprouts was significantly (*P* < 0.001) different from those of all five distinguished compartments of the seed tuber (Table S7). At phylum level, the bacterial community of the sprout was dominated by *Actinobacteria*, which were detected at a relative abundance of 72% of the total community, whereas *Firmicutes* (15%) and *Proteobacteria* (11%; Fig. S9) were also abundantly detected on sprouts. Also on sprouts, our analysis revealed a significant impact of plant genotype on microbial community composition (*P* = 0.001; Fig. 6a). Interestingly, 4 of the 6 varieties of sprouts emerging from seed tubers originating from different production fields had distinct microbiomes (Fig. 6b-g). These results indicate that the sprout-associated microbiome is influenced by plant genotype, but also by the field of production of the seed tuber.

**Fig. 6.**
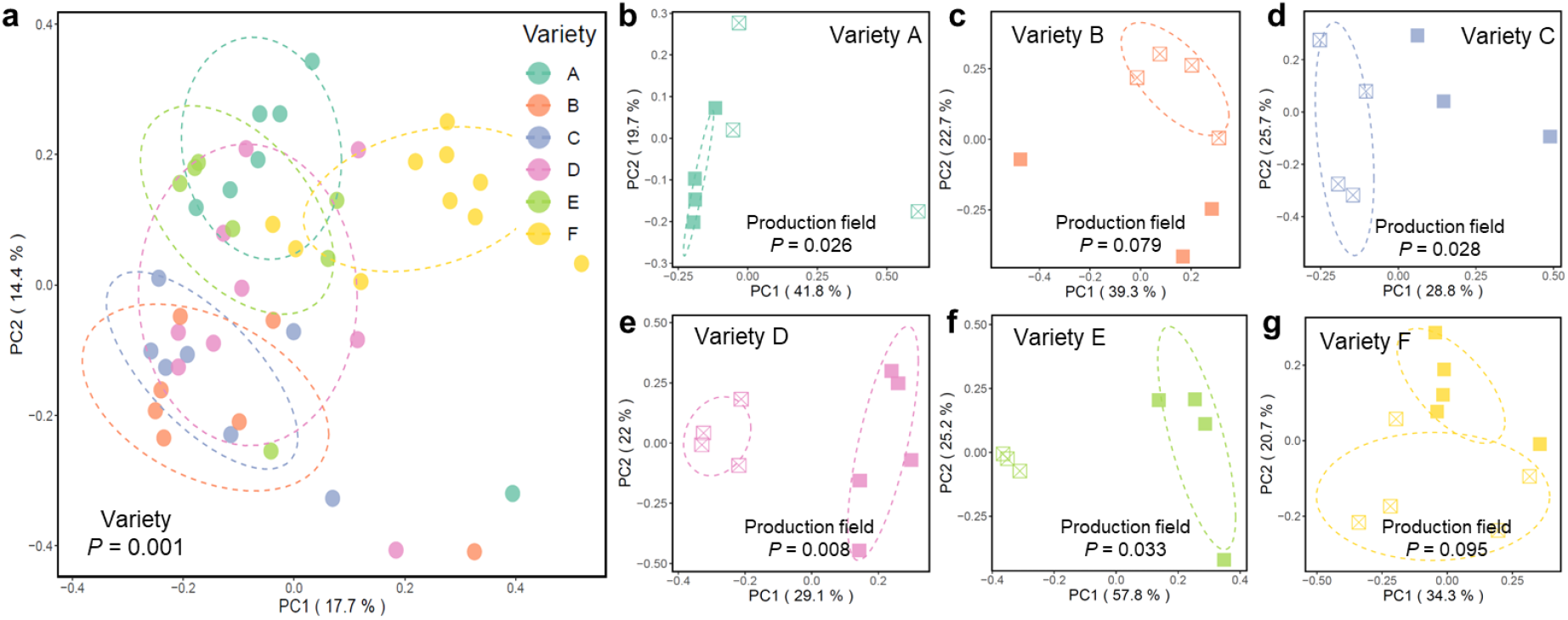
Field of production of the seed tuber affects the sprout microbiome. PCoA of bacterial sprout microbiomes of **a)** all varieties together and **b-e)** each variety separately. Each color represents one variety. Open and closed symbols represent distinct seed tuber production fields. The *P* from PERMANOVA is shown in each PCoA plot. Each sprout sample is a pool of 3-4 sprouts from one single tuber. Each ellipse represents a 68% confidence region and depicts the spread of data points within each group.

We next compared the microbiomes of the sprouts to the distinct compartments on the seed tubers that were analyzed above to identify the sources for the sprout microbiome. For bacteria, the analysis revealed that 79% (177 of 223) of the ASVs detected in the sprout microbiome were also detected in the microbiomes of at least one of the five seed tuber compartments (Fig. 7a). Thirty-one percent of these ASVs (70 of 223) were present in all compartments, but these 70 ASVs represented on average 60% of the total abundance of the sprout microbiome. Concomitantly, the 46 sprout-unique ASVs only made up 1.2% of the total bacterial abundance on the sprout (Fig. 7a). Thus, with 98.8% of the total bacterial abundance on the sprout, the seed tuber was the main source of the sprout microbiome in this soil-free system. Nonetheless, the taxonomic composition of the sprout microbiome was distinct from the compartments on the seed tuber (Fig. S9), indicating that the sprout compartment favors proliferation of a distinct subset of microbes that originate from the seed tubers.

We further analyzed whether specific compartments on the seed tuber contribute differentially to the sprout microbiome. Of the 223 bacterial ASVs detected on sprouts, 148 ASVs (66%) were also detected in adhering soil, 124 in heel end (56%), 128 in peel (57%), 109 in eye (49%) and 103 in flesh compartments (46%; Fig. 7b). We subsequently identified the top 18 most-abundant bacterial ASVs (ASVs with relative abundances over 1%) in the sprouts that made up 80% of the total bacterial sprout community and were able to trace them back in at least 2 of the 5 tuber compartments, but with significantly lower abundances comparing within the sprouts (Fig. 7c). For fungi, 8 ASVs out of the 74 ASVs that were detected in sprout samples were not found in any of the tuber compartments, and the 8 ASVs represented only 2% of the sprout fungal community (Fig. S10a). On the other hand, 46% (34 of 74) of the sprout ASVs were present in all compartments and represent on average up to 65% of the total abundance of the sprout fungal community (Fig. S10b). Furthermore, the top 16 most-abundant fungal ASVs (ASVs with relative abundances over 1%) in the sprout totaled 95% of the fungal sprout community (Fig. S10c). The relative abundance of these 16 fungal ASVs in the sprout did not differ significantly (ANOVA, Turkey, *P* > 0.05) between the distinct tuber compartments (Fig. S10c). Together these data show that both bacteria and fungi on seed tubers have the potential of being vertically transmitted to the sprouts, and that the sprout compartment subsequently promotes proliferation of a select number of microbes that are relatively lowly abundant in all compartments of the seed tubers.

**Fig. 7.**
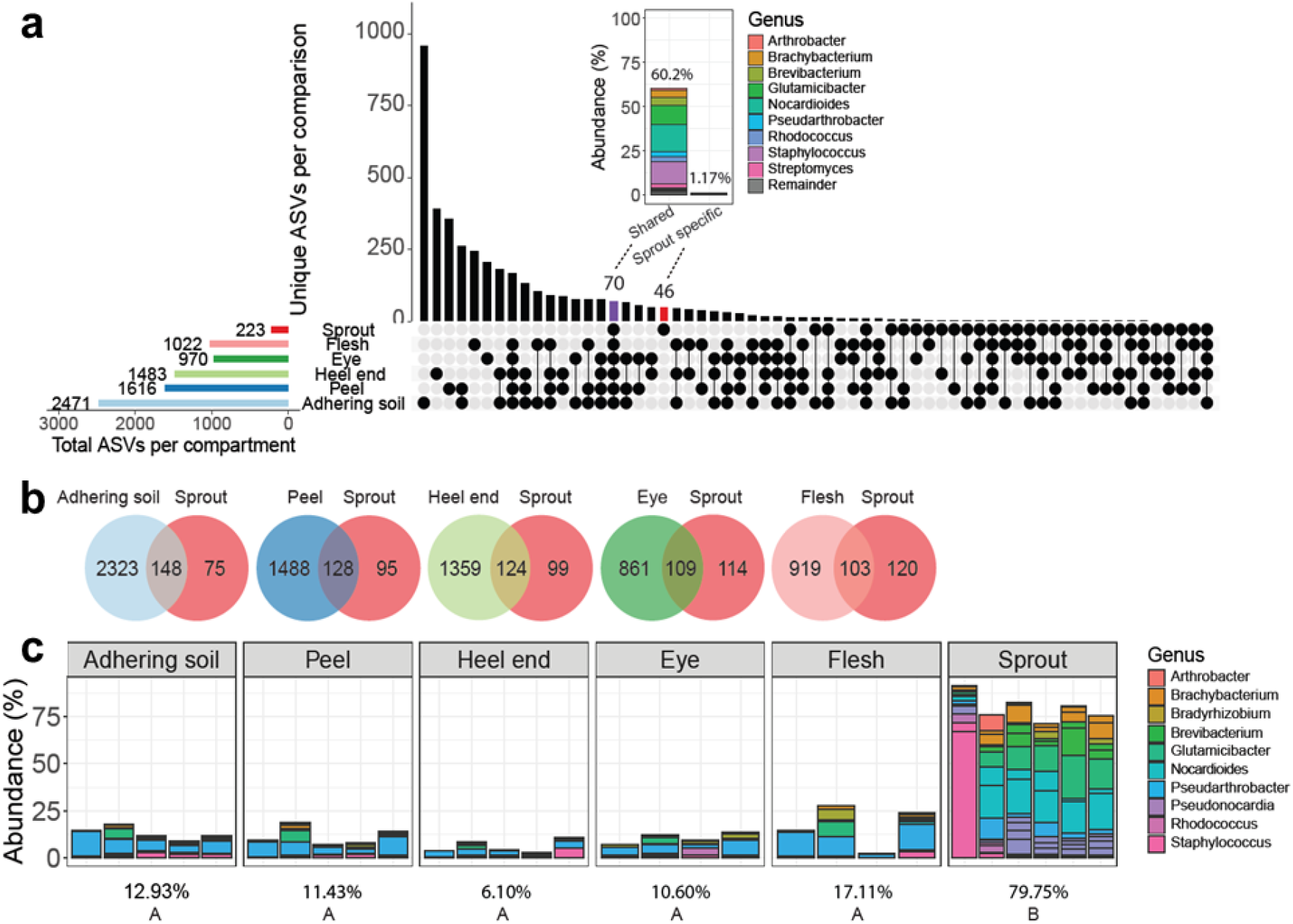
The sprout microbiome is derived from diverse seed tuber tissues. **a** UpSet plot shows shared and unique ASVs of each compartment of Variety A. Each row represents a sample type, and each column represents a set of ASVs, where filled-in black dots with an edge between the dots indicates that these ASVs are present in multiple sample types. The sets are ordered by the number of ASVs as indicated by the bar plot above each category. The total ASVs in each sample type is indicated by the rotated bar plot on the left. The inlay shows the abundance of ASVs (46) that are unique to sprouts and of sprout ASVs (70) that are shared with all tuber compartments. **b** Venn diagrams of ASVs shared between each tuber compartment and the sprout of Variety A. Color represents different compartment. **c** The distribution of the top 18 most-abundant sprout ASVs in all compartments of Variety A. Color represents the genus of the ASVs. The percentage under each figure shows the relative abundance of these top sprout ASVs in each compartment. Capital letters indicate significant difference (*P* < 0.05) in agglomerated abundance of the top sprout ASVs as determined by ANOVA with Tukey’s post-hoc test.

## Discussion

It has been well established that both soil type and plant genotype are important drivers in the assembly of plant-root associated microbial communities ^25^. However, seeds are also a source of microbiota that can be transmitted to the plants that develop from them ^26, 27^. Potatoes are vegetatively propagated by transplantation of relatively large seed tubers that contain a complex microbiome. Here we studied how the microbiome of seed potatoes is affected by the field of production and whether the seed tuber microbiome associated with production fields is transmitted to the emerging potato plant in the next season.

First, we analyzed two important factors that likely determine potato tuber microbiome composition. Soil has been reported to be the main source of microbes that colonize potato roots and tubers ^20, 28, 29^. In addition plant genotype is a factor that shapes plant-associated microbiomes^30^. Plant roots actively and dynamically secrete root exudates that can selectively promote or deter specific microbes ^31, 32^. Although up to 85% of the total dry matter produced by the potato plant can accumulate in the tubers ^33^, it is unclear whether the tuber actively exudes metabolites to interact with the microbiome. In this light, it has been reported that the tuber surface is low in nutrients and that the limited nutrients that are available to the microbiome are a result of cell decay or lesions only ^34^. Tubers might therefore control soil microbiota to a much smaller extent compared to roots. In line with this, Buchholz, Antonielli ^20^ and Nahar, Floc’h ^35^ reported that the microbiome found on potato tubers is largely independent from the potato genotype. Also, Weinert et al. show that tuber-associated bacteria were not strongly affected by the plant genotype although a few cultivar-dependent taxa were identified ^36, 37^.

In our study, however, when growing different genotypes in the same field we observed that not only root, but also the tuber-associated bacterial and fungal communities were significantly affected by the potato genotype (Fig. 3). Moreover, we found that the influence of potato genotype is larger on the tuber microbiome than on the potato root microbiome (Fig. 3, Table S1). This suggests that potato plants do exert control on the tuber microbiota, just like they selectively shape their root microbiomes. Nonetheless, up to half of the bacterial ASVs found on seed tubers harvested from one field were not found on seed tubers from the same variety that originated from other production fields (Fig. 1, Fig. S5). Field of production determined more than half of the bacterial variation of the seed tubers (Fig. 1, Table S1, S2). These results indicated that field of production is dominating over genotype and is playing an even more vital role in tuber-associated microbiome assembly than potato genotype, confirming previous findings ^20, 29, 35^.

Interestingly, we observed that both roots and daughter tubers in our trial field harbored microbiomes that were distinguishable by the production field of their seed tuber. This implies that there is intergenerational or vertical transmission of microbes from the seed tuber to the emerging plant and subsequently to the newly emerging tubers, the latter most likely via the stolon. In this light, Vannier et al. ^38^ reported that both bacteria and fungi of the clonal plant *Glechoma hederacea* can be transmitted to daughter plants through the stolon. In potato, some bacteria may migrate via the xylem or intracellular spaces to the above ground tissues of the potato plants as well as the stolon ^39^ and subsequently into the emerging tubers ^20^. These studies suggest that vertical transmission of microbes from one potato generation to the next is possible. In our study, we observe around a quarter of bacterial and up to half of fungal communities in the daughter tubers and roots overlapped with the seed tuber microbiomes (Fig. 4, Fig. S6). However, when we looked at ASVs that were uniquely found on roots and daughter tubers that originate from seed tubers from a specific production field, we see that a very small part of these ASVs (< 0.5%) is also detected uniquely on the seed tubers from that production field (Fig. 4, Fig. S6). We conclude that, based on the tractable vertical transmission of field-unique microbes, intergenerational transmission of microbiota is minimal and cannot explain the effects of field of production on microbiomes in the subsequent crop.

To better understand the early events in transmission of specific microbiome members from the seed tuber to plants emerging from these tubers, we analyzed the microbial composition of sprouts geminated in a soil-free system and compared it to the microbial communities of different compartments of the seed tubers. Firstly, we observed that the tuber’s adhering soil, peel, heel end, eye and flesh constitute distinct compartments that have significantly different microbiomes (Fig. 5, Fig. S7). Apparently the physical and chemical characteristics and activities in these distinct microhabitats ^40, 41^ select for different microbes. Moreover, the bacterial richness decreased from the surface of the tuber inwards (Fig. S8). Arguably this is a result of physical exclusion of microbes by the barrier function of the distinct tuber tissues and increased selective pressure inside the tuber by a combination of e.g., plant immunity and oxygen limitation ^42^.

In order to focus on the transmission from seed tuber to its sprouts without the interference of the soil, we subsequently analyzed the microbiomes of sprouts emerging from the seed tubers in a soil-free system. Our results showed that the early stage of microbial community assembly in the sprouts are genotype related. Moreover, sprouts emerging from tubers of the same genotype but originating from different production fields still show to some extent distinct microbial patterns (Fig. 6). These results indicate that the influence of tuber genotype and the field of seed tuber production can largely determine the early-stage microbial assembly on the potato sprouts. Moreover, the top 18 most abundant bacterial ASVs, comprising almost 80% of the total bacterial communities on the sprouts, could be traced back to the seed tuber compartments that we analyzed (Fig. 7, Fig. S10).

However, these sprout-abundant ASVs microbiome comprised a significantly smaller part of the total bacterial microbiome in the different seed tuber compartments. This suggests that the most abundant ASVs on the sprouts originate from diverse compartments of the seed tuber, and their proliferation was specifically stimulated by the sprout.

Together our data show that microbiome composition is intergenerationally affected by the field of production of the seed tuber. The potato tuber and root microbiomes on the daughter plants were comprised mostly of microbes derived from the soil environment in which the next-season potato plants were cultivated. The composition of a potato tuber microbiome is typically influenced by a combination of factors: the resident soil microbiome, potato genotype, and the specific physical, chemical, and (micro)biological conditions under which the tubers develop. In this study we demonstrate that the potato tuber microbiome is also affected by the field in which the seed tuber was produced. However, although we show that vertical transmission of microbes can occur from seed tuber to the emerging sprouts in a soil free system, most microbes that occur on the roots and daughter tubers of field-grown potato cannot be traced back to the population of seed tubers from which they emerged. We speculate that the abiotic and biotic environmental conditions in the fields of production differentially imprinted the seed tubers, leading to so far unknown epigenetic and/or metabolic changes in the seed tubers that in turn differentially altered interactions of the emerging plant with the soil microbiome, resulting in distinguishable microbiome signatures on daughter tubers and roots, depending on the field of production of the mother seed tuber.

In conclusion, we show that seed tuber imprinting by the field of production shapes the microbiome of the emerging potato plant. As it is accepted that plant microbiomes contribute to plant nutrition and health, the initial microbiome is a much-undervalued trait of seed tubers specifically, or planting materials in general. Elucidating the relative importance of the initial microbiome and the mechanisms by which the origin of planting materials affect microbiome assembly will pave the way for the development of microbiome-based predictive models that may predict the quality of seed tuber lots, ultimately facilitating microbiome-improved potato cultivation.

## Materials and Methods

### Potato varieties

In total, 5 potato varieties form the Royal HZPC Group and Averis Seeds B.V. were used in this study, namely variety *Colomba* (Variety A), *Innovator* (Variety B), *Festien* (Variety C), *Challenger* (Variety D), *Sagitta* (Variety E) and *Seresta* (Variety F).

### Sampling of seed tubers and post-storage seed tubers

In the autumn of 2018, seed tubers of two potato varieties (labelled A and B in this study to protect the commercial interests of the potato breeding companies that produced them) were harvested from 3 fields of production for Variety A and 3 other fields for Variety B (Fig. S1a-b). These tubers were shipped to a central location where they were subsequently stored in the dark at 4 °C. Seed tubers were taken from cold storage and sampled in December 2018 as “seed tuber” and July 2nd, 2019, as “post-storage seed tuber”. For seed tuber samples, peels were sampled from 24 seed tubers per production field and the peels of 6 tubers were pooled into a composite replicate sample, resulting in 4 replicated samples per variety per field. For post-storage seed tuber samples, peels were sampled from 36 seed tubers per field and the peels of 6 tubers were pooled into a composite replicate sample, resulting in 6 replicated samples per variety per field. In total, 144 seed tubers and 216 post-storage seed tubers were sampled and resulted in 24 seed tuber samples and 36 post-storage seed tuber samples. These samples were frozen in liquid N2, freeze-dried and stored in 50-mL falcon tubes at -20 °C prior to analysis.

### Sampling of daughter tubers and roots emerging from seed tubers

Seed tubers of Variety A and B of the above-mentioned 6 production fields were subsequently planted in a single trial field near Veenklooster (Fig. S1a; GPS location: 53.30353, 6.02670), the Netherlands. The chemical composition of this sandy field was analyzed by Normec Groen Agro Control B.V. and found to contain 1630 mg N/kg, 34 mg P2O5/l, 108 mg K/kg, 216 mg MgO/kg, 9 mg Na/kg, 3.4% organic matter and a sulfur supply capacity 7.2kg S/ha per year. The field pH was 5.1 and the cation exchange capacity was 57 mmol/kg. On April 16^th^, 2019, 24 seed tubers were planted in each of the 4 replicate plots which were randomly distributed across the field. On July 2^nd^, 2019, 4 potato plants were collected from the centre of each plot, from which the root material of each plant was sampled as a root sample, resulting in 4 root samples per plot. In detail, for each plant, the loosely attached soil was shaken off the roots, then the roots were cut into 5 cm fragments by sterile scissors and a random subset of the root fragments were stored in a 50-mL falcon tube. In the meantime, the peel of 6 newly formed tubers of each plant were samples and pooled as a composite daughter tuber sample, resulting in 4 daughter tuber samples per plot. For both tuber and root samples, the soil tightly attached to the peel and root was retained. In total, 96 potato plants and 576 daughter tubers were sampled resulting in 96 root and 96 daughter tuber samples. These samples were freeze-dried and stored in 50-mL falcon tubes at -20 °C prior to analysis.

### Sampling of seed tuber compartments

To dissect the contribution of microbiomes of different seed tuber compartments, namely peel, eyes, heel ends, flesh, and adhering soil (Fig. S1c), in shaping the sprout microbiome, we made use of material from a parallel study in which we harvested tubers from 6 potato varieties produced in 25 distinct production fields (Variety A from 5, Variety B from 5, Variety C from 3, Variety D from 5, Variety E from 5, and Variety F from 2 fields, respectively; Fig. S1). In detail, the adhering soil was gently rubbed from the tuber surface and collected in 50-mL falcon tubes. Subsequently, 1 cm thick cores were sampled from potato heel ends and eyes using a sterilized Ø 0.6-cm metal corer. Then, peel was sampled from around the minor axes of a tuber using a sterilized peeler. Flesh was sampled by halving a tuber using a sterile scalpel and sampling 1-cm core using a sterile Ø 0.6-cm metal corer from the centre of the tuber. Samples from 50 seed tubers were pooled into a single sample per compartment per field. In total, 1250 tubers were sampled to access the microbial composition of different tuber compartments, resulting in 125 compartment samples. These samples were freeze-dried and stored in 50-mL falcon tubes at -20 °C prior to analysis.

### Sampling of sprouts

To study early events in transmission of specific microbiome members from seed tubers to plants emerging from these tubers, the sprout microbiome was characterized. Seed tubers of all 6 varieties (Variety A-F) from 12 of the above mentioned 25 fields were employed to study the sprout microbiome (Fig. S1a-b). Five replicate tubers collected from each production field were germinated on sterile Petri dishes in dark conditions (20 °C and RH 68%). These 60 seed tubers were randomized in 6 trays and the position of the trays were rotated every day. After 7 days, 3−4 sprouts were removed from each tuber using a sterile scalpel and pooled as a composite sample. These 60 sprout samples were freeze-dried and stored in 2-mL Eppendorf tubes at -20 °C prior to analysis.

### Sample grinding

To grind the samples in high-throughput, four 5-mm sterile metal beads were added to freeze-dried samples in 50-mL falcon tubes and placed in a custom-made box. The samples were ground for 9 min on maximum intensity in a SK550 1.1 heavy-duty paint shaker (Fast & Fluid, Sassenheim, the Netherlands). Freeze-dried sprout samples were ground in 2-mL Eppendorf tubes with one 5-mm sterile metal bead per tube with a Tissuelyzer at 30 Hz for 1 min.

### DNA isolation, library preparation and sequencing

Genomic DNA was isolated from ±75 mg potato powder per sample using a Qiagen Powersoil KF kit. The KingFisher™ Flex Purification System machine was used for high throughput DNA isolation. DNA was quantified using a Qubit® Flex Fluorometer with the Qubit dsDNA BR Assay Kit (Invitrogen, Waltham, MA, USA) and normalized to a concentration of 5 ng/μl. The resulting DNA samples were then stored at -20 °C.

Bacterial 16S ribosomal RNA (rRNA) genes within the V3–V4 hypervariable regions were amplified using 2.5 μL DNA template, 12.5 μL KAPA HiFi HotStart ReadyMix (Roche Sequencing Solutions, Pleasanton, USA), 2 μM primers B341F (5’-TCGTCGGCAGCGTCAGATGTGTATAAGAGACAGCCTACGGGNGGCWGCAG-3’) and B806R (5’-GTCTCGTGGGCTCGGAGATGTGTATAAGAGACAGGACTACHVGGGTATCTAATCC-3’) ^43^ with Illumina adapter sequences in combination with 2.5 μM blocking primers mPNA (5’-GGCAAGTGTTCTTCGGA-3’) and pPNA (5’-GGCTCAACCCTGGACAG-3’) in 25 μL reactions. Blocking primers were used to avoid the amplification of mitochondrial (mPNA) or plastidial (pPNA) RNA from the plant host ^44^. Cycling conditions for 16S rRNA were (1) 95 °C for 3 min; (2) 95 °C × 30 s, 75 °C × 10 s, 55 °C × 30 s, 72 °C × 30 s, repeated 24 times; (3) 72 °C × 5 min; (4) hold at 10 °C.

Fungal internal transcribed spacer 2 (ITS2) DNA was amplified using 2.5 μL DNA template, 12.5 μL KAPA HiFi HotStart ReadyMix, 2 μM primers fITS7(5’-TCGTCGGCAGCGTCAGATGTGTATAAGAGACAGGTGARTCATCGAATCTTTG-3’) and ITS4-Rev (5’-GTCTCGTGGGCTCGGAGATGTGTATAAGAGACAGTCCTCCGCTTATTGATATGC-3’) with Illumina adapter sequences in combination with 2 μM blocking primers cl1ITS2-F (5’-CGTCTGCCTGGGTGTCACAAATCGTCGTCC-3’) and clITS2-R (5’-CCTGGTGTCGCTATATGGACTTTGGGTCAT-3’) in 25 μL reactions ^43^. Cycling conditions for ITS2 were (1) 95 °C for 3 min; (2) 95 °C × 30 s, 55 °C × 30 s, 72 °C × 30 s, repeated 9 times; (3) 72 °C × 5 min; (4) hold at 10 °C.

For both PCR reactions, DNA was cleaned using the KingFisher™ Flex Purification System. Twenty μL of vortexed AMPure XP Beads (Beckman Coulter, Brea, USA) were added to 25 μL of PCR product in a KingFisher™ 96 deep-well plate. Beads with adjoined DNA were washed by subsequent transfer to 3 KingFisher™ 96 deep-well plates with 80% ethanol and DNA was then eluted in 30 μL C6 elution buffer from the Qiagen Powersoil KF kit.

Index PCR reactions were performed using standard Illumina i7 (N701-N712) index primers for columns and Illumina i5 (N501-N508) index primers for rows of each plate. Five μL DNA sample was added to a mix of 2.5 μL 2 μM index primer, 12.5 μL KAPA HiFi HotStart ReadyMix and 5 μL Milli-Q H2O. Cycling conditions for index PCRs were (1) 95 °C for 3 min; (2) 95 °C × 30 s, 55 °C × 30 s, 72 °C × 30 s, repeated 9 times for 16S or 24 times for ITS2; (3) 72 °C × 5 min; (4) hold at 10 °C. After the index PCR, DNA was cleaned using the abovementioned cleaning protocol. DNA concentrations of all PCR products were measured using a Qubit® Flex Fluorometer with the Qubit dsDNA BR Assay Kit (Invitrogen, Waltham, MA, USA) and normalised to 2 ng/μL, after which the samples were pooled and sent for Illumina V3 2x300 bp MiSeq sequencing at USEQ (Utrecht, the Netherlands).

### Microbial community analysis and statistics

Both 16S and ITS2 rDNA raw sequencing reads were denoised, joined, delineated into amplicon sequence variants (ASVs), and assigned taxonomy in the Qiime2 (v.2019.7) environment ^45^. Datasets were demultiplexed and then filtered using the DADA2 pipeline ^46^. ASVs with less than 30 reads or present in less than 3 samples across all samples within a dataset were removed to minimize potential errors in sequencing. The representative sequences were subsequently taxonomically classified using a classifier trained with the 99% OTU threshold SILVA database ^47^ for bacteria and UNITE reference database (v.8.0)^48^ for fungi. For bacteria, we removed remaining 16S reads annotated as mitochondria or chloroplasts and kept only reads assigned to Bacteria. On average, the mitochondrial and chloroplast reads together accounted for 46%, 21%, 11% and 3% of 16S reads in the seed tuber, seed tuber after storage, daughter tuber and root samples, respectively, and 0.08%, 22%, 26%, 46%, 67% and 90% in the adhering soil, heel end, peel, eye, flesh and sprout, respectively. For fungi, we removed remaining ITS reads assigned as *Viridiplantae* and *Protista* and kept only reads assigned to Fungi. On average, plant-originated reads accounted for 53%, 16%, 78% and 30% of ITS reads in the seed tuber, seed tuber after storage, daughter tuber and root samples, respectively; and 2%, 11%, 27%, 35 %, 55% and 46% in the adhering soil, heel end, peel, eye, flesh and sprout, respectively.

The datasets with samples from seed tubers, post-storage seed tubers, daughter tubers, and root samples were rarefied to 10000 bacterial and 4000 fungal reads per sample, respectively. The datasets with samples from five compartments of the seed tuber were rarefied to 8000 reads per sample, for both bacterial and fungal reads.

Bray-Curtis dissimilarity matrices were created in QIIME2 and visualized in R using the Qiime2R and *ggplot2* package. Permutational multivariate analysis of variance (PERMANOVA, 999 permutations) tests were performed using QIIME2 to test the effect of different factors on the microbiome composition. Kruskal-Wallis tests were performed to test for differences in community diversity and evenness. Distance matrices were created separately for each generation and variety to compare the seedlots within the varieties using PERMANOVA tests. Venn diagrams were conducted by R package VennDiagram (v1.7.1, https://CRAN.R-project.org/package=VennDiagram). UpSet plots were generated by R package UpSetR ^49^. Sankey diagrams were produced by R package ggalluvial (http://corybrunson.github.io/ggalluvial/).

## Data availability

The datasets generated during and/or analyzed during the current study are available from the corresponding author on reasonable request.

## Code availability

Custom code for the analyses in the current study are available from the corresponding author on reasonable request.

## Acknowledgements

We gratefully acknowledge the valuable contributions of the Royal HZPC Group and Averis Seeds B.V. in providing the seed tuber material and supporting the field trial. Their collaboration was essential to the successful execution of this research. Special thanks are extended to Doretta Akkermans, Falko Hofstra and Martzen ten Klooster from HZPC Holding B.V. for their contribution to the sample collection. Additionally, we acknowledge the funding support received from Europees Landbouwfonds voor Plattelandsontwikkeling (ELFPO) on the “Flight-to-vitality” project, which greatly facilitated the completion of this study. This work was also partly supported by the Dutch Research Council (NWO) through the Gravitation program MiCRop (grant no. 024.004.014). The farm, potato and root icons used in this manuscript were designed using images from Flaticon.com. We sincerely appreciate the contributions and support from all individuals and organizations involved in making this research possible.

## Author contributions

R.L.B., P.A.H.M.B., R.J and C.M.J.P. conceived and designed the study. Y.S., C.D.J. and J.H.H.M. conducted the experiments and collected the data. Y.S. and J.S. performed the statistical analysis. Y.S. drafted the manuscript, and R.L.B., P.A.H.M.B., and C.M.J.P. provided critical revisions. All authors approved the final version of the manuscript.

