## Supplementary Figures for "Seed tuber imprinting shapes the next-generation potato microbiome"

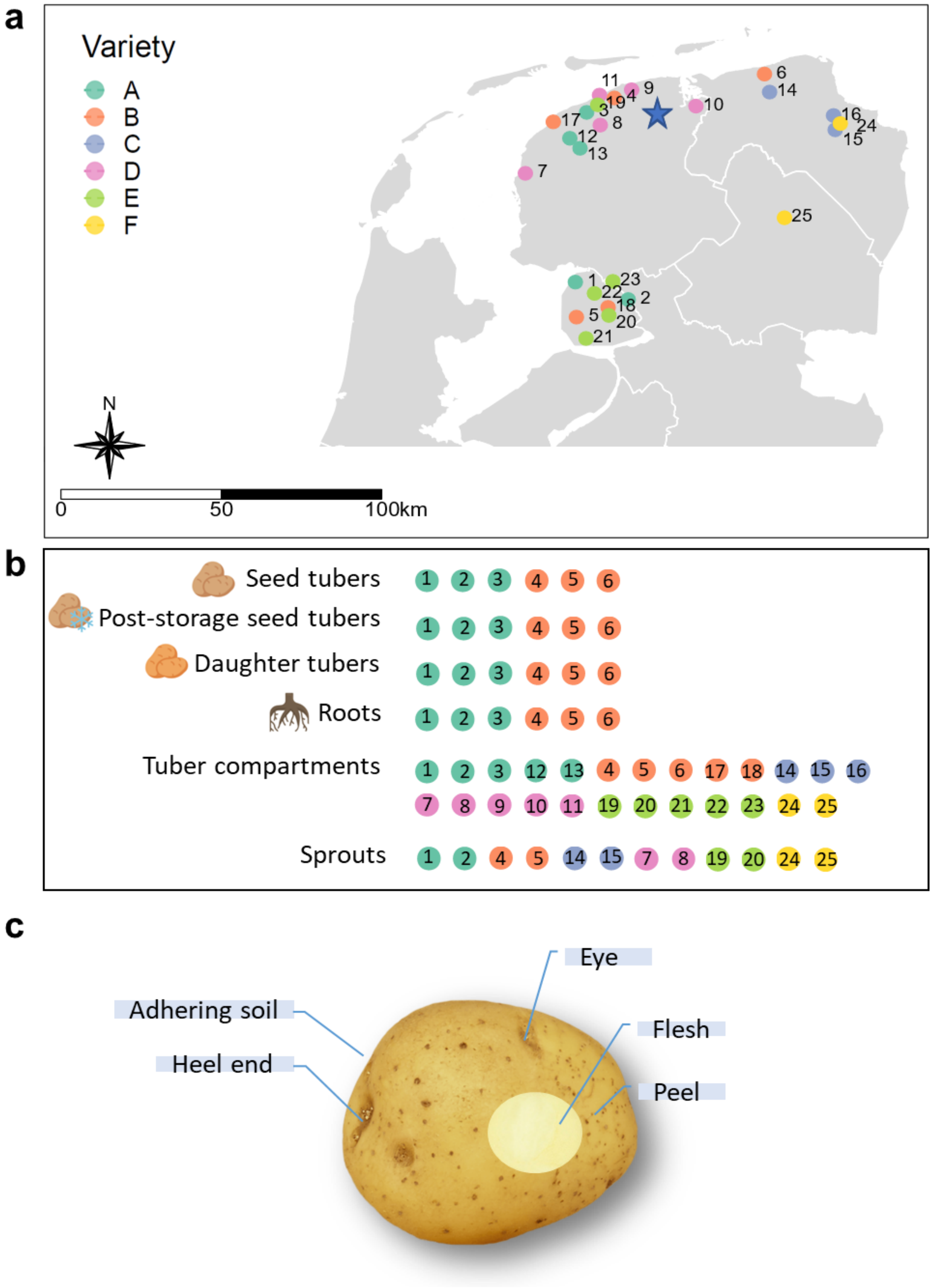


**Fig. S1 Experimental design.** **a** Map of the Netherlands with farms employed in this study. Color represents the variety of the seed tubers sampled from each field of production. All production fields are numbered from 1 to 25. The star represents the location of the field trial in Veenklooster. **b** Schematic diagram of fields involved in each data set. **c** Schematic diagram of five seed tuber compartments, adhering soil, potato heel end (where the tuber was generated from mother plant), potato eye (where the tuber will germinate and form daughter plant), potato peel and flesh.

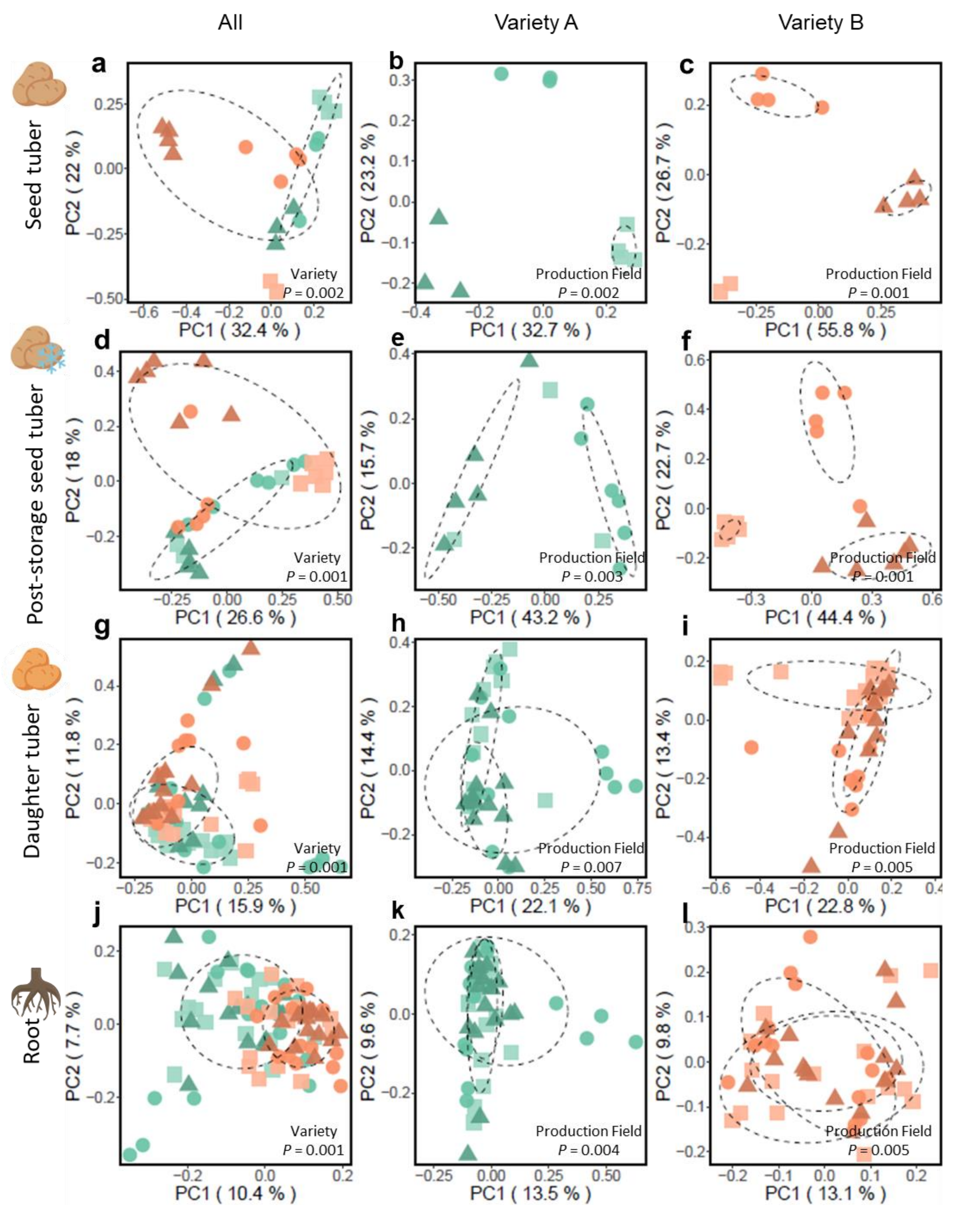


**Fig. S2 Fungal community composition of seed tubers, post-storage seed tubers, daughter tubers and roots.** Principle component analysis (PCoA) of ITS amplicon sequencing data of **a-c)** seed tubers of Variety A and/or B, **d-f)** post-storage seed tubers of Variety A and/or B, **g-i)** daughter tuber of Variety A and/or B, **j-l)** root of Variety A and/or B. Each symbol represents the bacterial community of one replicate potato peel sample. Each seed tuber sample consists of a pool of potato peels collected from 6 seed tubers. Each daughter tuber sample consisted of a pool of potato peels collected from 6 daughter tubers of one plant. Each root sample is a subset of the whole root of the same plant from which the daughter tubers were sampled. For each variety, 4 replicate samples were collected from each of the 3 fields of production. Green symbols represent Variety A and orange symbols represent Variety B. Different shapes within a same color represent distinct production fields. The *P*-value from PERMANOVA is shown in each PCoA plot. Each ellipse represents a 68% confidence region and depicts the spread of data points within each group.

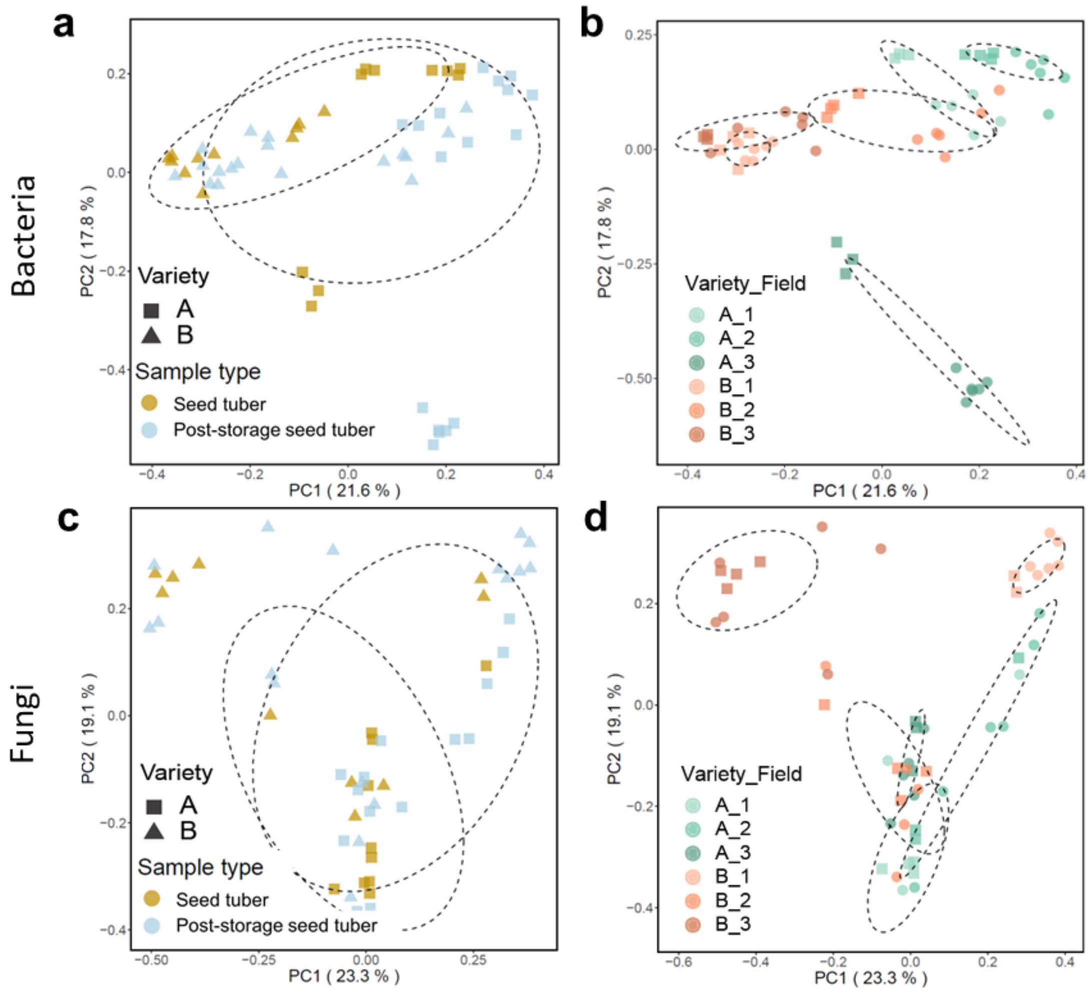


**Fig. S3 Bacterial and fungal community of seed tubers and post-storage seed tubers.** PCoA of bacterial communities of seed tubers and post-storage seed tubers, colored by **a)** sample type and **b)** production field, respectively. PCoA of fungal communities of seed tubers and post-storage seed tubers, colored by **c)** sample type and **d)** production field, respectively. Square symbols represent Variety A and triangle symbols represent Variety B. Each ellipse represents a 68% confidence region and depicts the spread of data points within each group. Each seed tuber sample is a pool of peel collected from 6 seed tubers.

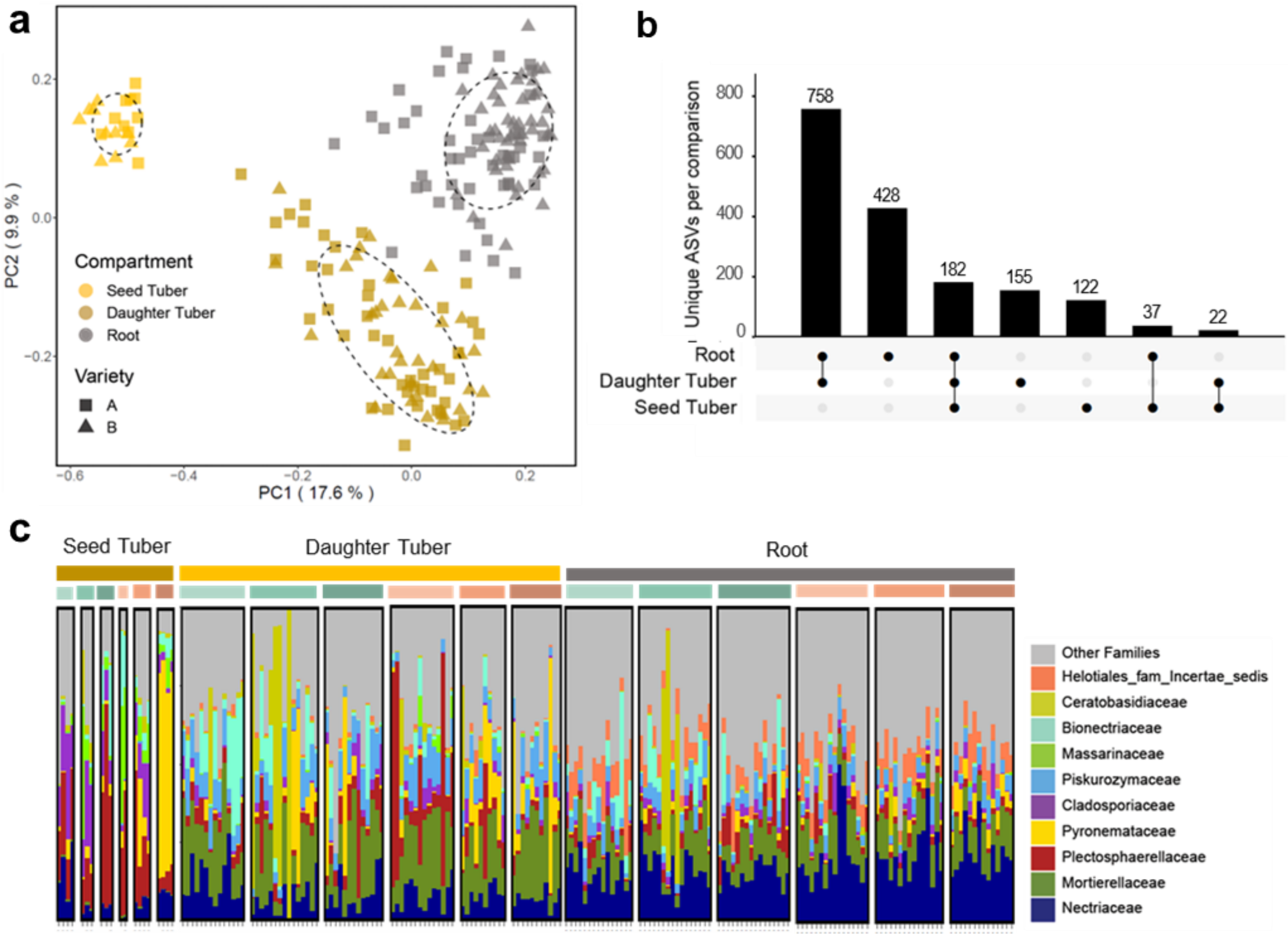


**Fig. S4 Analysis of fungal communities on seed tubers from different production fields and their roots and daughter tubers in a next-season trial field. a** PCoA of potato-associated fungal communities of seed tubers, daughter tubers and roots. Square symbols represent Variety A and triangle symbols represent Variety B. Colors represent different sample types. Each ellipse represents a 68% confidence region and depicts the spread of data points within each group. **b** UpSet plot with fungal ASVs present in seed tubers, daughter tubers and roots from all varieties. Each row represents a sample type, and each column represents a set of ASVs, where filled-in black dots with an edge between the dots indicates that these ASVs are present in multiple sample types. The sets are ordered by the number of ASVs as indicated by the bar plot above each category. **c** Fungal community phylogenetic composition (relative abundance) of different sample types at family level. Each stacked column represents an independent sample. Different colors within a column represent different families. Only the top 10 most abundant family were colored individually, all the rest are colored in gray as the "Other Families". Samples are clustered by sample type and field of production, which is shown by colored bar on top of the column. Each seed tuber sample is a pool of peel collected from 6 seed tubers. The root material of each plant was sampled as a root sample, the peel of 6 newly formed tubers of each plant were samples and pooled as a composite daughter tuber sample.

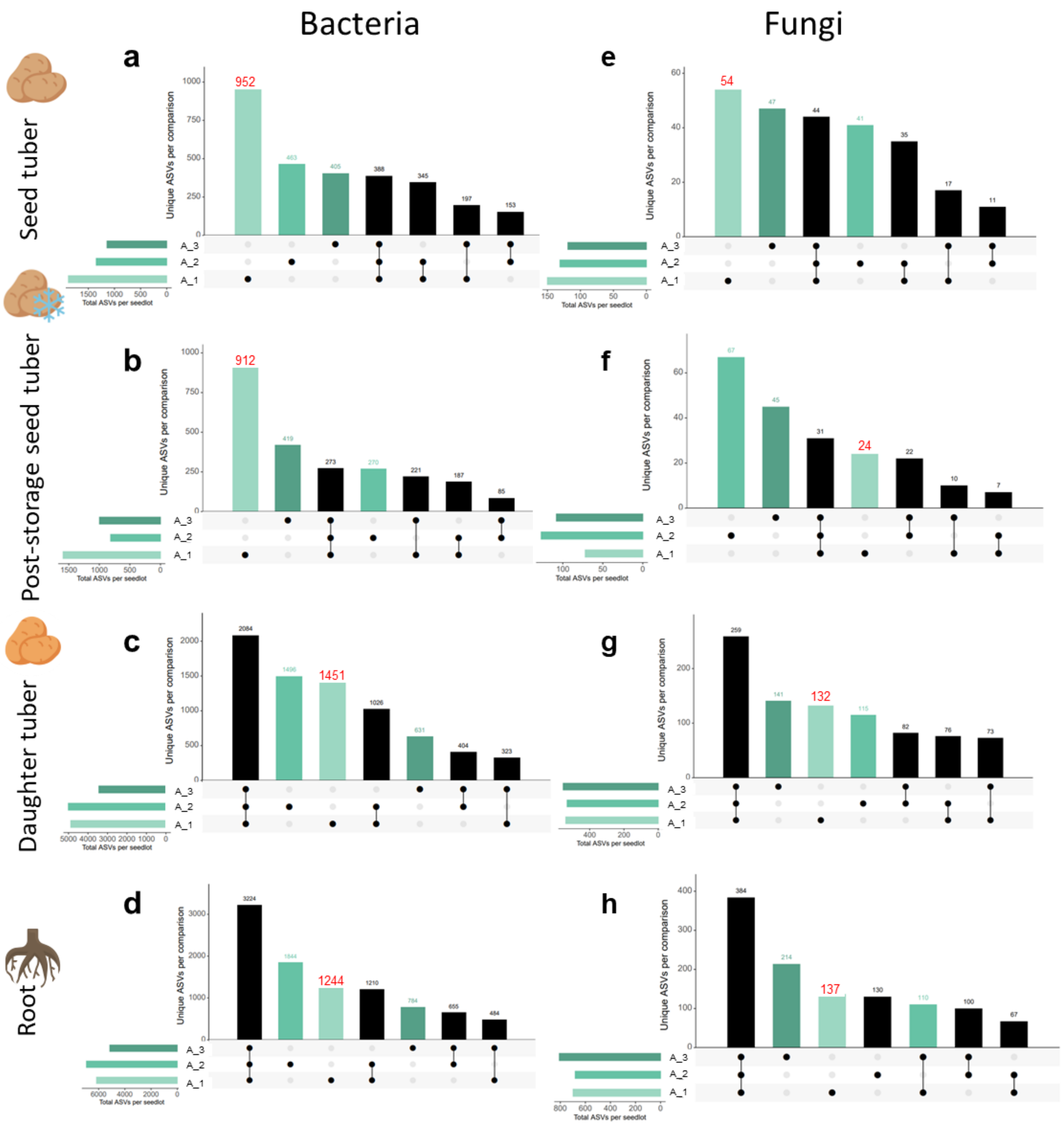


**Fig. S5 UpSet plot with ASVs present in each production field of Variety A in seed tubers, post-storage seed tubers, daughter tubers and roots.** Each row represents samples from one field of production, and each column represents a set of ASVs, where filled-in black dots with an edge between the dots indicate that these ASVs are present in multiple sample types. The sets are ordered by the number of ASVs as indicated by the bar plot above each category. The total ASVs in each production field are indicated by the rotated bar plot on the left. The sets of ASVs used in Fig. 4 and Fig. S6 are highlighted in red in the Upset plots.

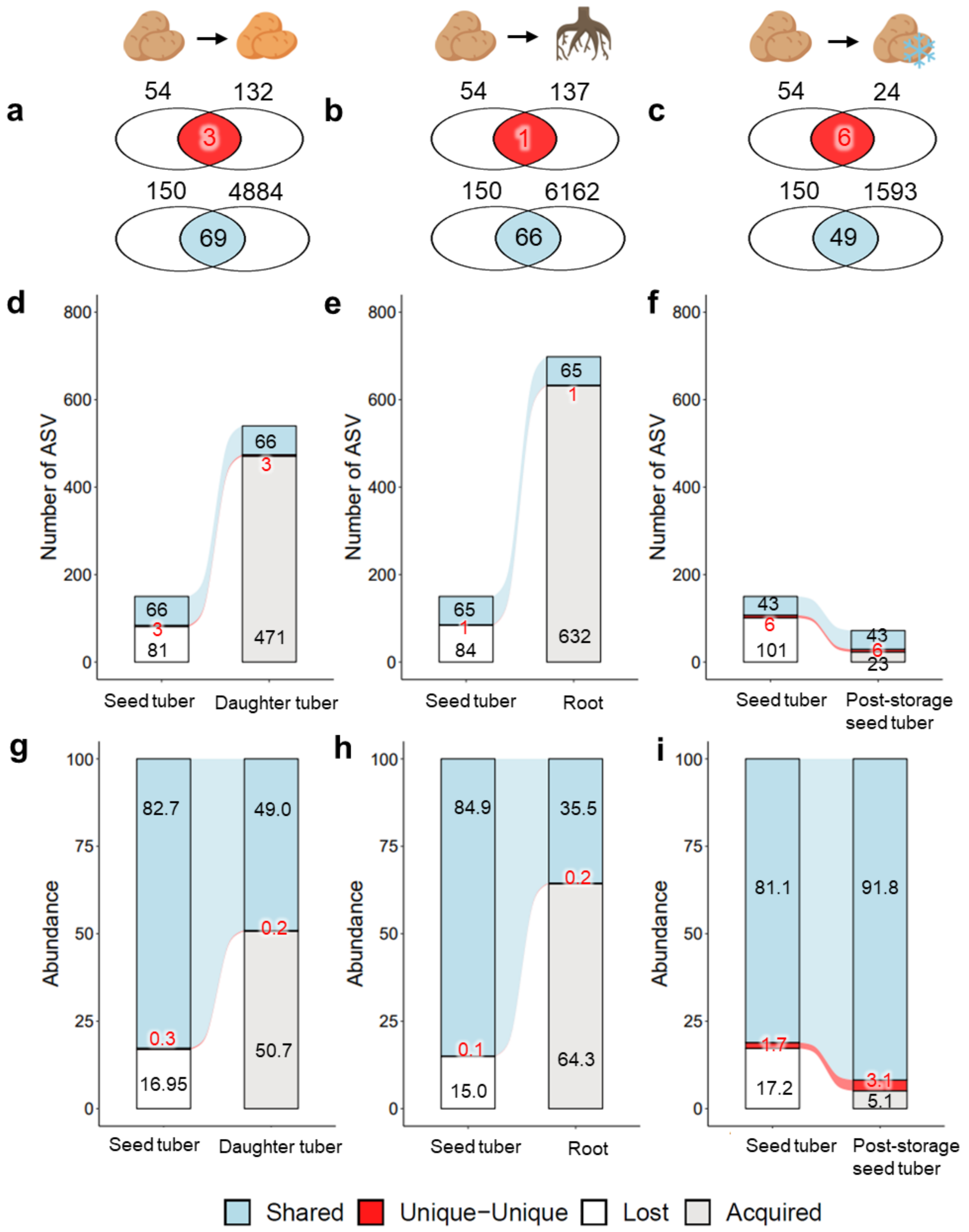


**Fig. S6 Comparison of fungal ASVs on daughter tubers, roots, post-storage seed tubers and seed tubers of Variety A originating from Field 1.** Venn diagrams showing the overlap between **a)** seed tubers and daughter tubers, **b)** seed tubers and roots, **c)** seed tubers and post-storage seed tubers of Field-1-unique fungal ASVs (in red) or all fungal ASVs (in blue). Sankey diagram of fungal ASVs transferred from seed tubers to **d, g)** daughter tubers and **e, h)** roots that emerged from the seed tubers; and **f, i)** post-storage seed tubers. "Shared" in blue represents ASVs detected on both sample types. "Unique-Unique" in red represents the overlap of Field-1-unique ASVs on both sample types. "Lost" in white represents ASVs lost from the seed tuber during vertical transmission. "Acquired" in light grey represents ASVs not transmitted from seed tubers but acquired from the environment. In **a-f),** numbers in the bars indicate the number of ASVs in each category mentioned above. In **g-i),** numbers in the bars indicate the accumulative relative abundance of ASVs in each category mentioned above.

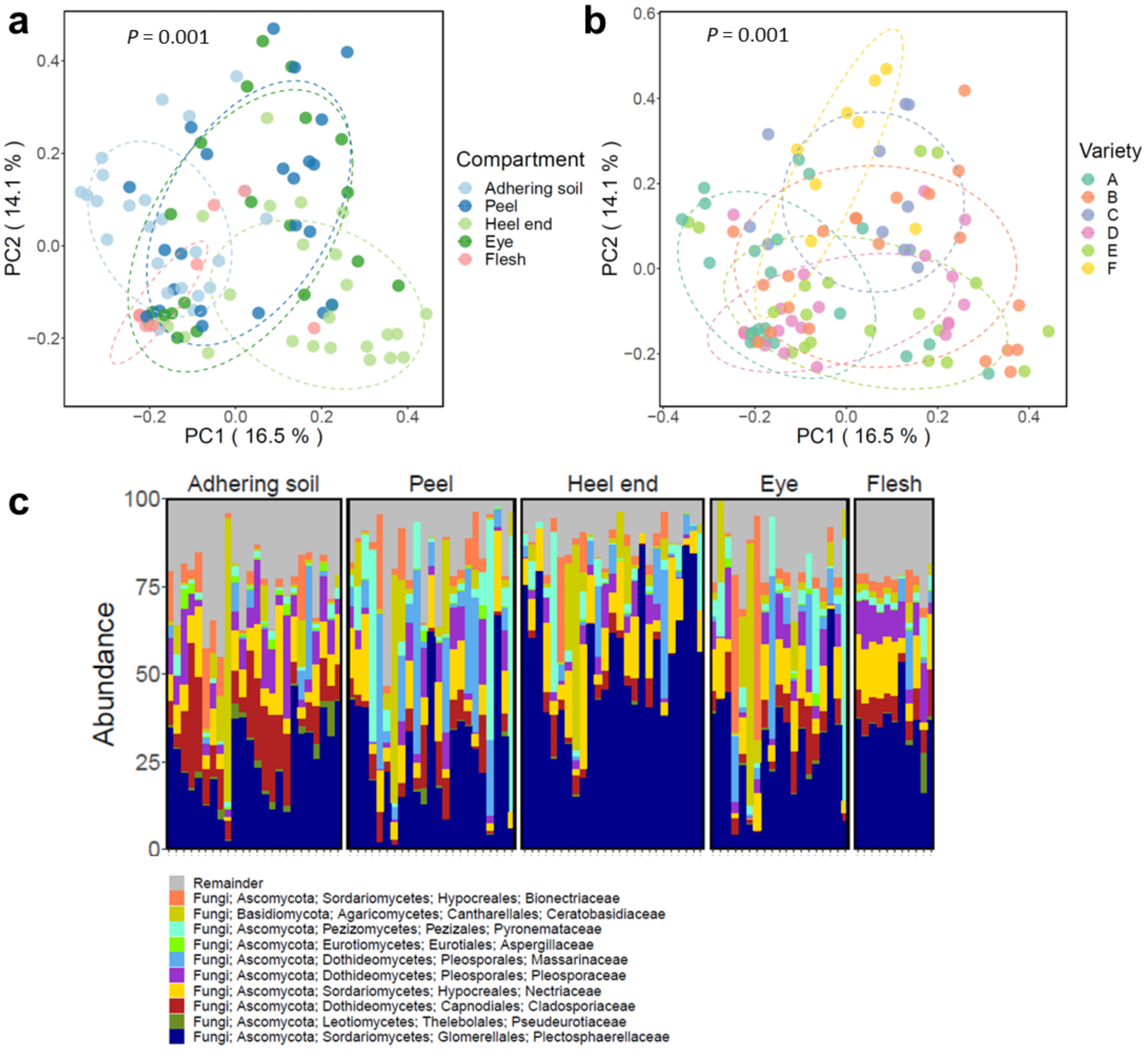


**Fig. S7 Different tuber compartments, namely potato flesh, heel end, peel, eye and adhering soil harbor distinct fungal communities.** PCoA of the potato tuber-associated fungal community based on ITS amplicon sequencing and colored by **a)** potato variety and **b)** potato compartment. The *P*-value from PERMANOVA is shown in each PCoA plot. Each ellipse represents a 68% confidence region and depicts the spread of data points within each group. **c** Bar plot shows the phylogenetic composition of fungal community. Only the top 10 most abundant families are colored individually, the other families are shown together in grey.

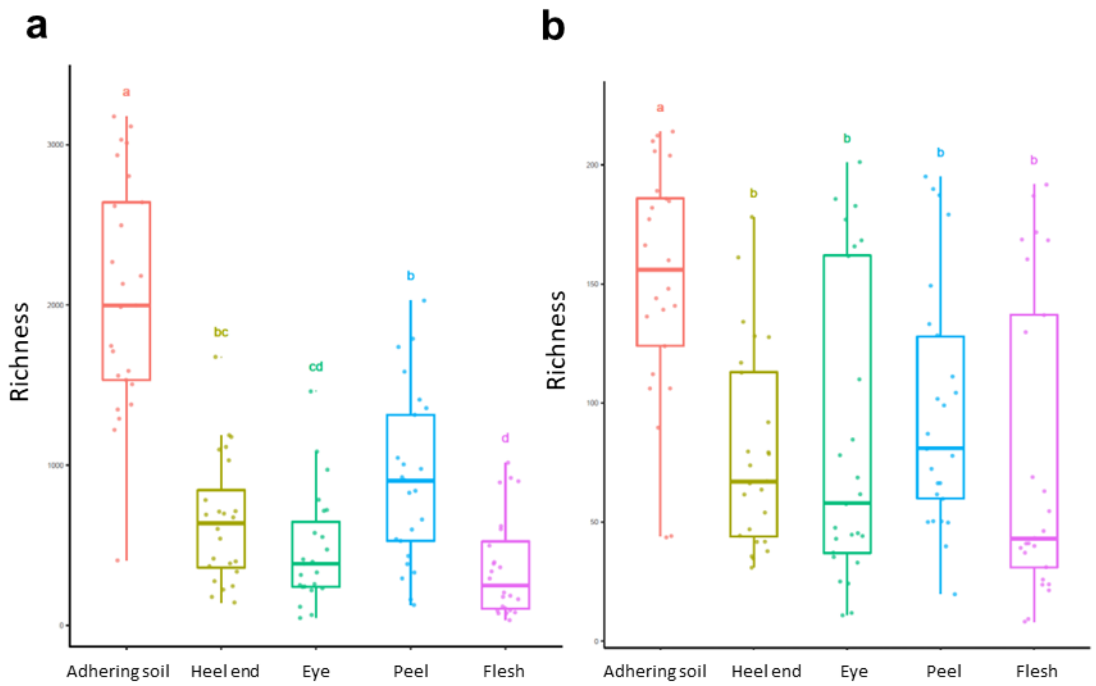


**Fig. S8** The Richness of **a)** bacteria and **b)** fungi community of 5 different tuber compartments.

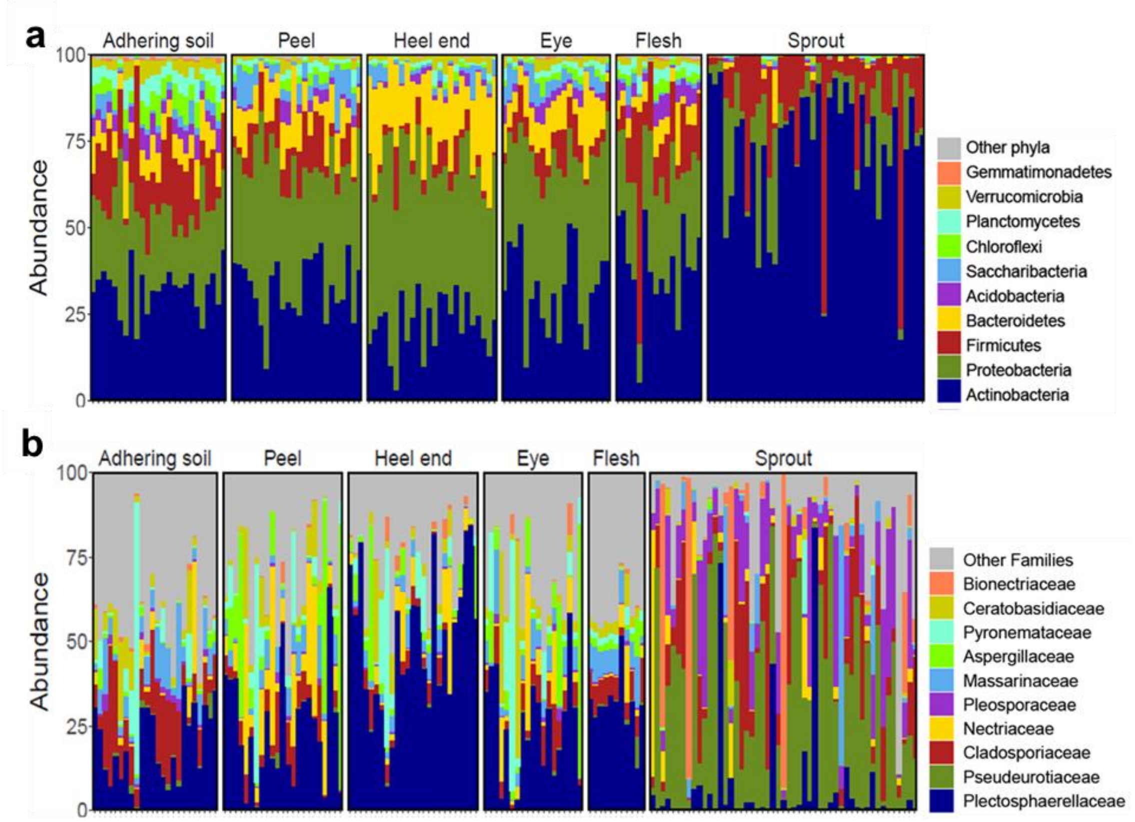


**Fig. S9 Sprout harbors distinct a) bacterial and b) fungal communities comparing with different tuber compartments, namely potato flesh, heel end, peel, eye and adhering soil.** The bar plot shows the phylogenetic composition. Only the top 10 most abundant **a)** phyla or **b)** families are colored individually, the rest are combined as other phyla or families.

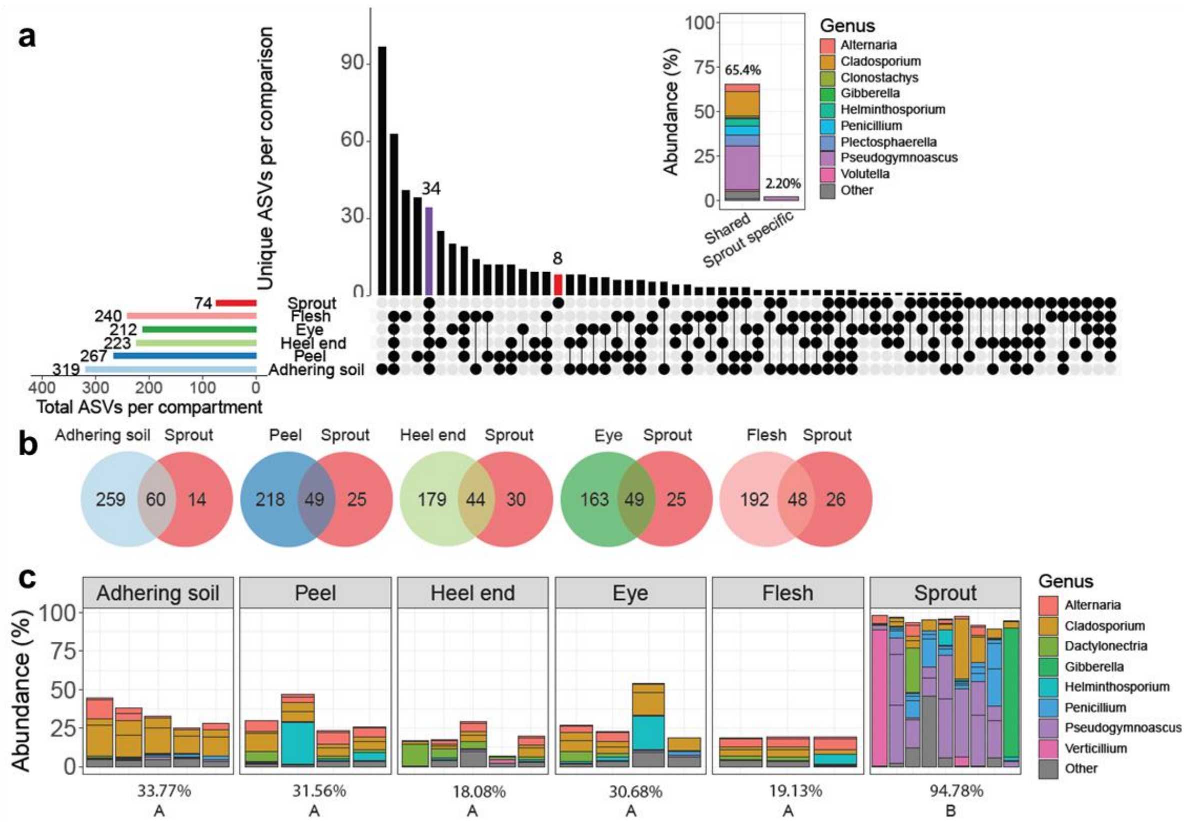


**Fig. S10 Selective assembly of the potato sprout fungal community. a** Upset plot shows shared and unique ASVs of each compartment of Variety A. Each row represents a sample type, and each column represents a set of ASVs, where filled-in black dots with an edge between the dots indicates that these ASVs are present in multiple sample types. The sets are ordered by the number of ASVs as indicated by the bar plot above each category. The total ASVs in each sample type is indicated by the rotated bar plot on the left. The inlay shows the abundance of ASVs (8) that are unique to sprouts and of ASVs (34) that are shared with all tuber compartments. **b** Venn diagrams of ASVs shared between each tuber compartment and the sprout of Variety A. Color represents different compartment. **c** The distribution of the sprout top 14 most- abundant ASVs in all compartments of Variety A. Color represents the genus of the ASVs. The percentage under each figure shows the relative abundance of these top sprout ASVs in each compartment. Capital letters indicate significant difference ($P < 0.05$) in agglomerated abundance of the top sprout ASVs as determined by ANOVA with Tukey's post-hoc test.
