## Supplementary Tables for "Seed tuber imprinting shapes the next-generation potato microbiome"

**Table S1** Results of Permutational multivariate analysis (PERMANOVA) for variance of the bacterial composition for seed tuber, post-storage seed tuber, daughter tuber and root samples at ASV level. Tests are based on Bray-Curtis dissimilarity distances and 999 permutations.

| Sample Type | Factors | Degrees of Freedom | Sums of Squares | Mean Squares | F.Model | $R^2$ | *P* |
|---|---|---|---|---|---|---|---|
| All Samples | Generation | 1 | 17.33 | 17.33 | 99.47 | 0.25 | 0.001 |
| | Sample Type | 2 | 10.65 | 5.32 | 30.55 | 0.15 | 0.001 |
| | Residuals | 239 | 41.63 | 0.17 | NA | 0.60 | NA |
| | Total | 242 | 69.60 | NA | NA | 1.00 | NA |
| Seed Tuber | Variety | 1 | 0.77 | 0.77 | 14.09 | 0.18 | 0.001 |
| | Production Field | 4 | 2.79 | 0.70 | 12.78 | 0.64 | 0.001 |
| | Residuals | 15 | 0.82 | 0.05 | NA | 0.19 | NA |
| | Total | 20 | 4.38 | NA | NA | 1.00 | NA |
| Post-storage Seed Tuber | Variety | 1 | 1.40 | 1.40 | 19.93 | 0.17 | 0.001 |
| | Production Field | 4 | 4.65 | 1.16 | 16.55 | 0.57 | 0.001 |
| | Residuals | 29 | 2.04 | 0.07 | NA | 0.25 | NA |
| | Total | 34 | 8.08 | NA | NA | 1.00 | NA |
| Daughter Tuber | Variety | 1 | 1.54 | 1.54 | 8.67 | 0.08 | 0.001 |
| | Production Field | 4 | 1.32 | 0.33 | 1.86 | 0.07 | 0.001 |
| | Residuals | 89 | 15.85 | 0.18 | NA | 0.85 | NA |
| | Total | 94 | 18.72 | NA | NA | 1.00 | NA |
| Root | Variety | 1 | 0.36 | 0.36 | 3.36 | 0.03 | 0.001 |
| | Production Field | 4 | 0.82 | 0.20 | 1.88 | 0.08 | 0.001 |
| | Residuals | 86 | 9.34 | 0.11 | NA | 0.89 | NA |
| | Total | 91 | 10.52 | NA | NA | 1.00 | NA |

**Table S2** Results of PERMANOVA for variance of the fungal composition for seed tuber, post-storage seed tuber, daughter tuber and root samples at ASV level. Tests are based on Bray-Curtis dissimilarity distances and 999 permutations.

| Sample Type | Factors | Degrees of Freedom | Sums of Squares | Mean Squares | F.Model | $R^2$ | *P* |
|---|---|---|---|---|---|---|---|
| All Samples | Generation | 1 | 14.58 | 14.58 | 72.43 | 0.23 | 0.001 |
| | Sample Type | 2 | 5.25 | 2.63 | 13.04 | 0.08 | 0.001 |
| | Residuals | 216 | 43.48 | 0.20 | NA | 0.69 | NA |
| | Total | 219 | 63.32 | NA | NA | 1.00 | NA |
| Seed Tuber | Variety | 1 | 0.71 | 0.71 | 8.28 | 0.17 | 0.001 |
| | Production Field | 4 | 2.33 | 0.58 | 6.85 | 0.55 | 0.001 |
| | Residuals | 14 | 1.19 | 0.09 | NA | 0.28 | NA |
| | Total | 19 | 4.23 | NA | NA | 1.00 | NA |
| Post-storage Seed Tuber | Variety | 1 | 0.77 | 0.77 | 5.33 | 0.10 | 0.001 |
| | Production Field | 4 | 3.67 | 0.92 | 6.34 | 0.46 | 0.001 |
| | Residuals | 25 | 3.61 | 0.14 | NA | 0.45 | NA |
| | Total | 30 | 8.05 | NA | NA | 1.00 | NA |
| Daughter Tuber | Variety | 1 | 0.60 | 0.60 | 2.85 | 0.03 | 0.001 |
| | Production Field | 4 | 1.66 | 0.42 | 1.97 | 0.10 | 0.001 |
| | Residuals | 71 | 15.01 | 0.21 | NA | 0.87 | NA |
| | Total | 76 | 17.28 | NA | NA | 1.00 | NA |
| Root | Variety | 1 | 0.72 | 0.72 | 5.01 | 0.05 | 0.001 |
| | Production Field | 4 | 0.97 | 0.24 | 1.69 | 0.07 | 0.001 |
| | Residuals | 86 | 12.35 | 0.14 | NA | 0.88 | NA |
| | Total | 91 | 14.04 | NA | NA | 1.00 | NA |

**Table S3** Results of PERMANOVA test of bacterial and fungal composition of seed tuber and post-storage seed tuber samples at ASV level. Tests are based on Bray-Curtis dissimilarity distances and 999 permutations with FDR correction for multiple comparisons.

| | Sample Type 1 | Sample Type 2 | Sample Size | Permuations | pseudo-F | p-value | q-value |
|---|---|---|---|---|---|---|---|
| Bacteria | Seed Tuber | Post-storage Seed Tuber | 56 | 999 | 4.223 | 0.001 | 0.001 |
| Fungi | Seed Tuber | Post-storage Seed Tuber | 51 | 999 | 2.345 | 0.019 | 0.019 |

**Table S4** Results of PERMANOVA test of bacterial and fungal composition of different sample types at ASV level. Tests are based on Bray-Curtis dissimilarity distances and 999 permutations with FDR correction for multiple comparisons.

| | Sample Type 1 | Sample Type 2 | Sample Size | Permuations | pseudo-F | p-value | q-value |
|---|---|---|---|---|---|---|---|
| Bacteria | Seed Tuber | Daughter Tuber | 116 | 999 | 38.908 | 0.001 | 0.001 |
| | Seed Tuber | Root | 113 | 999 | 59.571 | 0.001 | 0.001 |
| | Daughter Tuber | Root | 187 | 999 | 61.369 | 0.001 | 0.001 |
| Fungi | Seed Tuber | Daughter Tuber | 97 | 999 | 24.680 | 0.001 | 0.001 |
| | Seed Tuber | Root | 112 | 999 | 44.832 | 0.001 | 0.001 |
| | Daughter Tuber | Root | 169 | 999 | 24.923 | 0.001 | 0.001 |

**Table S5** Results of pairwise Adonis test of bacterial composition of 5 tuber compartments at ASV level. Tests are based on Bray-Curtis dissimilarity distances and 999 permutations with FDR correction for multiple comparisons.

| Group 1 | Group 2 | Sample size | Permutations | pseudo-F | p-value | q-value |
|---|---|---|---|---|---|---|
| Adhering soil | Heel end | 45 | 999 | 7.926 | 0.001 | **0.002** |
| Adhering soil | Eye | 36 | 999 | 4.250 | 0.001 | **0.002** |
| Adhering soil | Peel | 47 | 999 | 4.183 | 0.001 | **0.002** |
| Adhering soil | Flesh | 34 | 999 | 2.210 | 0.002 | **0.003** |
| Heel end | Eye | 31 | 999 | 1.322 | 0.055 | 0.061 |
| Heel end | Peel | 42 | 999 | 2.479 | 0.001 | **0.002** |
| Heel end | Flesh | 29 | 999 | 3.091 | 0.001 | **0.002** |
| Eye | Peel | 33 | 999 | 1.243 | 0.143 | 0.143 |
| Eye | Flesh | 20 | 999 | 1.592 | 0.012 | **0.015** |
| Peel | Flesh | 31 | 999 | 1.546 | 0.009 | **0.013** |

**Table S6** Results of pairwise Adonis test of fungal composition of 5 tuber compartments at ASV level. Tests are based on Bray-Curtis dissimilarity distances and 999 permutations with FDR correction for multiple comparisons.

| Group 1 | Group 2 | Sample size | Permutations | pseudo-F | p-value | q-value |
|---|---|---|---|---|---|---|
| Adhering soil | Eye | 44 | 999 | 3.409 | 0.001 | **0.002** |
| Adhering soil | Flesh | 35 | 999 | 2.849 | 0.003 | **0.004** |
| Adhering soil | Heel end | 49 | 999 | 10.021 | 0.001 | **0.002** |
| Adhering soil | Peel | 47 | 999 | 4.386 | 0.001 | **0.002** |
| Eye | Flesh | 31 | 999 | 1.929 | 0.030 | **0.033** |
| Eye | Heel end | 45 | 999 | 2.976 | 0.002 | **0.003** |
| Eye | Peel | 43 | 999 | 0.614 | 0.830 | 0.830 |
| Flesh | Heel end | 36 | 999 | 5.041 | 0.001 | **0.002** |
| Flesh | Peel | 34 | 999 | 2.584 | 0.011 | **0.014** |
| Heel end | Peel | 48 | 999 | 4.351 | 0.001 | **0.002** |

**Table S7** Results of pairwise Adonis test of bacterial and fungal composition of sprout and 5 tuber compartments at ASV level. Tests are based on Bray-Curtis dissimilarity distances and 999 permutations with FDR correction for multiple comparisons.

| | Sample Type 1 | Sample Type 2 | Sample Size | Permuations | pseudo-F | p-value | q-value |
|---|---|---|---|---|---|---|---|
| Bacteria | Adhering soil | Sprout | 65 | 999 | 23.955 | 0.001 | 0.001 |
| | Eye | Sprout | 60 | 999 | 19.134 | 0.001 | 0.001 |
| | Flesh | Sprout | 56 | 999 | 13.605 | 0.001 | 0.001 |
| | Heel end | Sprout | 64 | 999 | 23.609 | 0.001 | 0.001 |
| | Peel | Sprout | 64 | 999 | 21.230 | 0.001 | 0.001 |
| Fungi | Adhering soil | sprout | 75 | 999 | 19.990 | 0.001 | 0.002 |
| | Eye | sprout | 70 | 999 | 14.987 | 0.001 | 0.002 |
| | Flesh | sprout | 62 | 999 | 13.753 | 0.001 | 0.002 |
| | Heel end | sprout | 76 | 999 | 25.635 | 0.001 | 0.002 |
| | Peel | sprout | 74 | 999 | 17.470 | 0.001 | 0.002 |
